# Integrative Transcriptomic Analysis of Anterior and Posterior IOP-Controlling Tissues in Glaucoma Reveals Enrichment of MHC-II Pathway and T-Cell Infiltration Signatures

**DOI:** 10.1101/2023.09.26.558362

**Authors:** Shuxin Liang, Shichen Zhang, Hongqiu Wang, Shaojun Tang

**Author notes:** The authors wish it to be known that the first two authors should be regarded as co-first authors.

## Abstract

**Background:** Glaucoma is a multifactorial neurodegenerative disease indicating by abnormalities in the whole eye, apart from optic neuropathy, and is often associated with abnormal intraocular pressure (IOP). However, the underlying pathological mechanisms of glaucoma and the management of IOP remain largely unaddressed. Emerging evidence suggests the involvement of immune changes in glaucoma. Therefore, characterizations of immune alternations by molecular profiling of IOP-controlling tissues from both anterior and posterior eye segments including trabecular meshwork (TM) and choroid may provide insights to the disease mechanism.

**Methods:** Bulk TM microarray data (GSE138125 and GSE27276) from primary open-angle glaucoma (POAG) and choroid single-cell transcriptome data (GSE203499) from glaucoma were downloaded from Gene Expression Omnibus (GEO) database. The two qualified TM datasets were integrated. Analyses of differential expression genes (DEGs), functional and pathway annotations, immune infiltration, and disease-related modules were performed using the POAG bulk data. Similarly, DEGs and pathway annotations, single-cell trajectory and switch gene analysis, and cell-cell communications were conducted using the choroid single-cell RNA-sequencing data. Finally, integrated analyses from the two studies were conducted to determine the glaucomatous changes between anterior and posterior IOP-controlling eye tissues.

**Results:** A total of 102 DEGs between POAG and healthy tissues were identified from the TM data. Gene set enrichment analysis revealed five glaucoma-enriched pathways: ATP generation and metabolism, cornification and keratinization, metabolism of reactive oxygen species (ROS), humoral immune response, and platelet aggregation. Significant immune infiltrations were observed in POAG tissue, including total T cells (excluding CD8^+^ T cells), NK cells, monocytic lineage, endothelial cells, and fibroblasts. Furthermore, the POAG enriched pathways are primarily from T-cell infiltrations. On the other hand, fourteen distinct cell clusters were identified from the choroid single-cell data. The most evident findings were fibrosis, represented by extracellular matrix (ECM), the actin-binding pathways enrichment in pericyte-fibroblast transition, reduction in light sensitivity of melanocytes, and complex immune changes among all cell types in the pathological conditions compared to control. Taken together, an integrated analyses between the TM microarray and choroid single-cell data result in 111 significantly dysregulated genes in the disease state. These key genes participated in immune and inflammatory reactions (DUSP1, ADM, CD74, CEBPD, HLA-DPA1, ZNF331), served as potential biomarkers for neurodegenerative and autoimmune diseases (such as NR4A2), or regulated membrane integrity, vasculature calcification, ECM interactions, and cell morphology (PTP4A1, MGP, and RASD1). Overall, our integrated data analyses might provide insight to the understanding of the disease mechanism.

**Conclusion:** By integrating bulk microarray data from TM and single-cell transcriptome data from the choroid, we systematically evaluated the glaucomatous transcriptomic changes between the anterior and posterior IOP-controlling tissue from ocular. Results indicated there was a significantly enrichment of genes belonging to the MHC-II pathway and T-cell infiltration in the disease state, opening new avenues for biomarkers discovery and therapeutic interventions to glaucoma.

## Background

Glaucoma, a neurodegenerative eye disease, stands as the leading cause of irreversible blindness worldwide, with primary open-angle glaucoma (POAG) being the most prevalent subtype ^1,2^. Despite extensive research efforts dedicated to understanding this disease, its underlying pathological mechanisms remain elusive, impeding the development of effective strategies for early screening, diagnosis, and treatment ^3^. Notably, glaucoma is recognized as a multifactorial condition, with pathological changes detectable throughout the entire eye, beyond optic neuropathy ^4^. This suggests that glaucomatous pathologies manifest across various structures, ranging from the anterior chamber and trabecular meshwork to the choroid, retina, and optic nerves ^2,4–6^. Among the multiple factors contributing to glaucoma, elevated or fluctuating intraocular pressure (IOP) emerges as the most significant characteristic in the majority of POAG cases, surpassing factors such as age, race, and family history ^7^. Excessive production or impaired drainage of aqueous humor (AH) in the anterior chamber of the eye serves as the primary cause of abnormal IOP ^8^.

The trabecular meshwork (TM), situated in the anterior segment of the eye, serves as the primary pathway for AH outflow, distinct from the uveoscleral outflow route, commonly referred to as the conventional pathway. In the context of POAG, where there are no obstructions narrowing or closing the anterior chamber angle, the TM plays a crucial role ^9,10^. Extensive evidence supports the presence of pathological changes within the TM in POAG, including increased resistance, thickening, and a reduction in TM surface area. However, the molecular mechanisms underlying these abnormalities are not yet fully understood ^11–14^.

As depicted in **Fig. 1**, the human TM (hTM) is composed of three distinct regions: the corneoscleral, uveal, and juxtacanalicular portions, extending from the iris to the Schlemm’s canal ^15^. The TM cells (TMCs) in the first two regions establish direct communication with the AH, forming a single layer and resting on a basement membrane. These cells cover an area constructed by collagen and elastin fibers and exhibit characteristics of both endothelial cells and macrophages. On the other hand, TMCs in the juxtacanalicular portion are dispersed within the extracellular matrix (ECM) and exhibit fibroblast and smooth muscle cell-like properties ^11,16^. Therefore, it is highly plausible that the pathological changes in the TM and their impact are multifaceted. Notably, the endothelial cell and macrophage characteristics of TMCs suggest their potential involvement in regulating oxidative stress and immune mediation.

**Figure 1:**
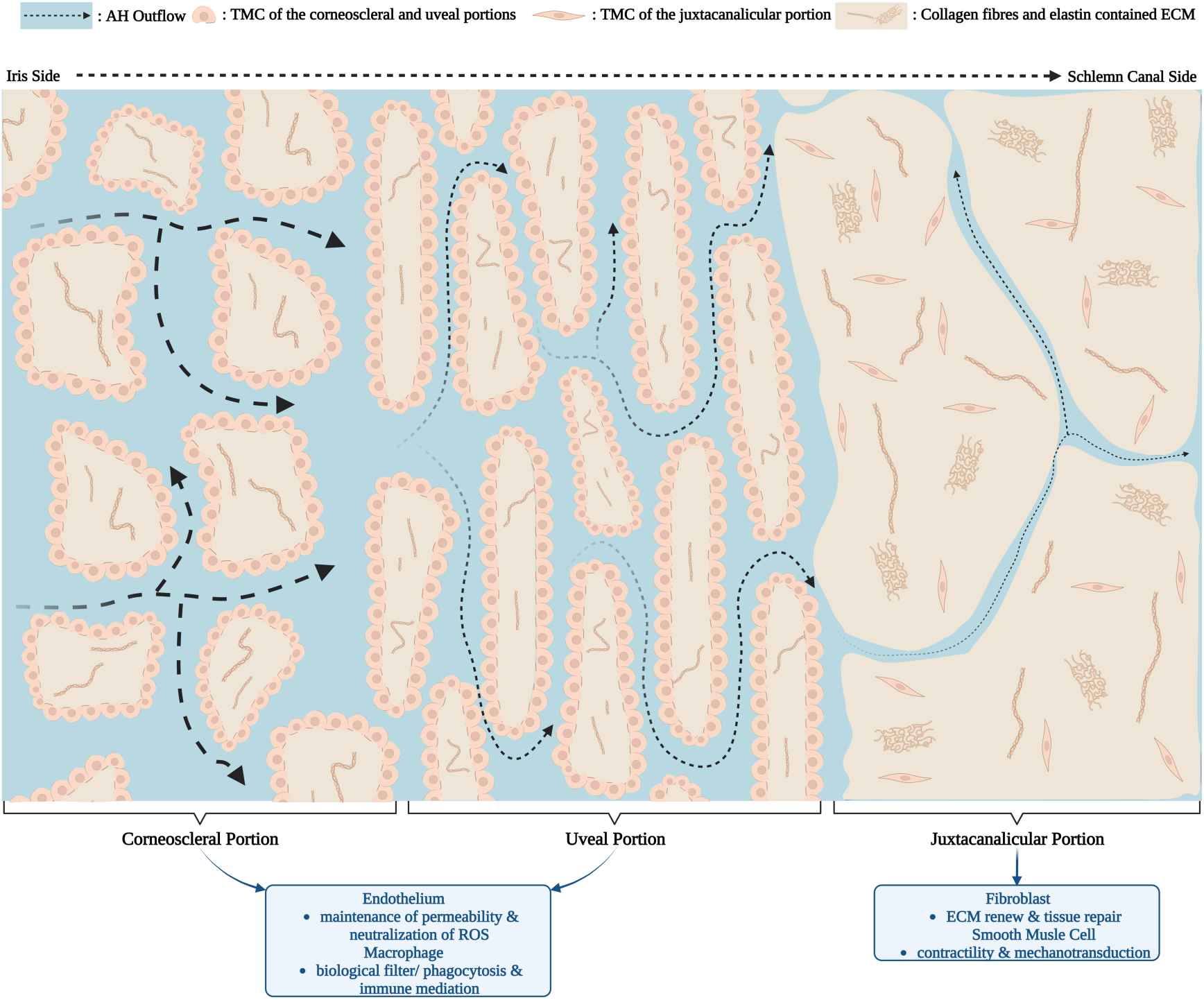
Schematic Representation of the Microscopic Structure of Normal Human Trabecular Meshwork (TM).

Apart from the TM, it is important to acknowledge the significance of the choroid, which contribute to the uveoscleral outflow of the AH. The choroid is a vascular tissue layer that located between the retina and the sclera. It serves as a supplier of oxygen and nutrients to the outer layer of the retina and plays a crucial role in regulating IOP ^17–20^. Similar to other blood vessels, the choroid is immunocompetent and composed of endothelial cells, pericytes, fibroblasts, melanocytes, and macrophages ^17^. As the nourishing tissue of the posterior segment of the ocular adjacent to the blood-retina barrier, it has an important role in glaucoma.

The immune aspects of glaucoma research deserve attention. The ocular system is immunoprivileged and maintains a state of immunosuppression. However, the influence of immune activation or disorders in glaucoma cannot be overlooked ^21,22^. Immunotherapy holds promise for the management and treatment of glaucoma. Evidence suggests the presence of activated innate and adaptive immune responses in glaucoma ^22^. For instance, glial cells such as microglia, astrocytes, and Müller cells, which contribute to immune surveillance in the retina, have been found to be activated in the early stages of glaucoma ^23^. Moreover, a correlation has been observed between elevated IOP and T cell-mediated autoimmunity, particularly in relation to HSP-specific responses ^24^. However, the specific HSP antigen and its interaction with distinct subtypes of T cells, leading to dominant roles in cross-immune reactions, remain unknown ^25^. Furthermore, the evidence is limited to the presence of autoantibodies in peripheral blood and relative changes in the neural retina. Given the immunomodulatory capabilities of the TM and choroid, it is worth investigating their roles in glaucoma, particularly in relation to immune-related changes.

In this study, our focus was on investigating gene expression patterns in the TM of patients with POAG. To ensure robust results and analyze a larger sample, we selected and integrated two qualified microarray datasets from the Gene Expression Omnibus (GEO) database. We employed gene set enrichment analysis (GSEA) and identified differentially expressed genes (DEGs) between the TM of POAG patients and control subjects. To gain further insights into the biological changes associated with POAG, we performed Gene Ontology (GO) annotations and Kyoto Encyclopedia of Genes and Genomes (KEGG) pathway analysis on the DEGs, confirming the presence of abnormalities in the identified genes.

To explore potential hub genes related to POAG, we constructed a protein-protein interaction (PPI) network using the Search Tool for the Retrieval of Interacting Genes (STRING) database and Cytoscape. Through this analysis, we identified thirty suspected hub genes. Additionally, we performed immune cell type deconvolution analysis to study the infiltration of primary immune cells, endothelium, and fibroblasts. The results of this analysis were incorporated as clinical traits in weighted correlation network analysis (WGCNA) to identify POAG-related gene modules and their association with immune changes.

To compare changes in the anterior and posterior segments of the glaucomatous eye, we analyzed single-cell transcriptome data from the choroid of patients with glaucoma, which has not been published previously. We performed DEGs analysis, pathway annotations, trajectory and switch gene analyses, and assessed cell communication. By comparing these findings with the results obtained from the integrated TM datasets, we aimed to obtain a comprehensive understanding of tissue abnormalities related to AH drainage and IOP regulation. Our analyses also shed light on the role of these abnormalities in immune disorders and the pathology of glaucoma.

## Methods

### TM datasets

#### Dataset selection and data preprocessing

Data from the GEO (https://www.ncbi.nlm.nih.gov/geo/)^26^ consisting of TM subjects with and without POAG, was downloaded. The search using keywords: primary open-angle glaucoma, POAG, trabecular meshwork, TM, Homo sapienS, and human, yielded three datasets: GSE138125, GSE27276, and GSE4316. However, GSE4316 was excluded due to its small sample size and suspiciously low-quality data ^27^.

After removing the lncRNA probes in GSE138125 and transforming the gene IDs, the mRNA data from the two log2-transformed datasets were merged into a single file. Only reads that were detected in both datasets were retained. Then, lowly expressed genes were filtered based on their counts per million values, ensuring that they were greater than 1 in over half of the samples.

To address batch effects, Combat normalization was performed using the SVA package ^28^ in R (Version 4.1.1). Finally, the distribution of samples based on gene expression profiles from different groups and datasets was visualized through principal component analysis (PCA) using the R packages FactoMineR ^29^ and factoextra ^30^ .

#### Differential expression and pathway enrichment analysis

The limma R package ^31^ was employed to identify DEGs between POAG and control (normal) participants. DEGs were determined based on the criteria of |log_2_FC| > 1 with an adjusted P value (*adj.P*) < 0.05. Subsequently, a volcano plot was generated using the ggplot2 package ^32^, and a heatmap illustrating all the DEGs was created using the pheatmap package ^33^. To gain a deeper understanding of the biological functions associated with the DEGs, a hierarchical clustering of genes into three levels was performed for GO-based biological processes (BP) annotation analysis. Additionally, both GO ^34^ term and KEGG ^35^ pathway analyses were conducted using the clusterProfiler R package ^36^ to provide further pathway information related to the DEGs. The obtained information was visualized using the enrichplot ^37^ and ggplot2 R packages. Furthermore, GSEA was conducted using the clusterProfiler R package with a significance threshold of *P.Val* < 0.05, and the results were visualized using the enrichplot R package.

#### STRING and PPI network analysis and hub gene identification

Protein-protein interaction (PPI) networks of the DEGs were constructed using the STRING database (https://string-db.org/) ^38^. In the PPI network, each pair of interactors among the DEGs possessed a combined confidence score of at least 0.4. To identify hub genes within the PPI networks, the cytoNCA ^39^ and CytoHubba ^40^ plugins of the Cytoscape software (version 3.9.1) ^41^ were utilized to analyze the level of connectivity. Based on the Maximal Clique Centrality (MCC) scores ^40^, the top 30 potential hub genes were selected, and the PPI networks were visualized using the Cytoscape software. Furthermore, the gene expression profiles of the 30 potential hub genes were plotted using the ggplot2 package after undergoing the Wilcoxon test. To evaluate the predictive efficiency for POAG, receiver operating characteristic (ROC) analysis was performed using the pROC R package ^42^ .

#### Immune cell type deconvolution analysis

The MCP-counter R package, which quantifies the abundance of distinct immune cells utilizing specific molecular markers ^43^, was employed to assess immune infiltration in each sample. Moreover, Wilcoxon tests were conducted to compare the infiltration levels of individual immune and stromal cell types between the POAG and control groups.

#### WGCNA analysis

A gene co-expression network was constructed using the WGCNA R package ^44^. Initially, the pickSoftThreshold function was utilized to determine the optimal power value for aligning the gene distribution with a connection-based scale-free network. Subsequently, the adjacencies were transformed into a topological overlap matrix (TOM), enabling the grouping of genes with similar expression patterns into distinct modules. A minimum genome of 30 was employed for the gene tree, and a tangent of 0.25 was utilized for the module tree, resulting in the formation of diverse modules. Modules exhibiting similar gene distribution were merged to form new modules.

Following the WGCNA analysis, a correlation analysis was conducted between the outcomes of the immune infiltration analysis and the co-expression modules enriched by WGCNA. Consequently, it was observed that the two most POAG-related modules exhibited a strong correlation with immune infiltration. Subsequently, the module eigengene-based connectivity was calculated to identify hub genes for further investigation. Hub genes were selected based on an absolute eigengene-based connectivities (kME) value higher than 0.8 for the two modules.

#### Functional analysis of POAG-related module genes from WGCNA

The PPI networks of the hub genes of the two modules were constructed using STRING and high-resolution bitmaps were downloaded (each pair of interactors in the hub genes had a combined confidence score of no less than 0.4). GO analysis was performed using R software to study the main biological functions of the genes.

### Choroid Dataset

#### Dataset selection and data preprocessing

Single-cell RNA sequencing data were obtained from the GEO dataset GSE203499, which does not have any previously reported articles associated with it. The dataset comprised of 11 patients with varying degrees of age-related macular degeneration (AMD), along with individuals suffering from cataracts, glaucoma, or hypertension. Choroidal samples were collected from patients diagnosed with early AMD and glaucoma, as well as from a control group with early AMD and a normal phenotype. This sampling strategy was employed to minimize the confounding effects of early AMD disease.

To analyze the data at the single-cell level, we utilized R (version 4.1.2) and Seurat (version 4.1.0) software ^45^. We initially applied the emptyDrop and PercentageFeatureSet functions to remove empty droplets and calculate mitochondrial gene percentages, respectively. Subsequently, we filtered out cells with feature counts > 4600, < 200, and > 25% mitochondrial transcripts to eliminate the effects of empty droplets, multiple droplets, and dead cells. Following this filtering step, we retained 27,686 cells containing 31,679 genes for downstream analysis.

For cell cycle analysis, we selected genes from the GO:0007049, GO:0044843, and GO:1902749 pathways, and then scored the cell cycles using the CellCycleScoring function. Next, log normalization was performed, and highly variable expression genes were selected using the VariableFeatures function for PCA downscaling. The top ten principal components were identified and used for clustering with a resolution of 0.6, determined via the ScoreJackStraw function and elbow plot. To visualize the results, we employed t-SNE and UMAP visualization techniques. Furthermore, specific marker genes in each cell cluster were identified using the FindAllMarkers function in the Seurat package.. Subsequently, we classified all choroidal cells into 14 distinct cell types using the marker genes provided in the cellmarker2.0 database.

#### Differential gene expression and pathway enrichment analyses

To identify DEGs between patients with and without glaucoma, we performed differential gene expression analysis using the FindMarkers function in the Seurat package. DEGs between normal and glaucomatous tissues were selected based on an *adj.P* < 0.05 and logFC > 0.58. Furthermore, we also identified DEGs in specific cell types. The up- and down-regulated DEGs were subjected to GO, KEGG, and Reactome ^46^ pathway enrichment analyses using the clusterProfiler package. Pathways with a *P.Val* < 0.05 were considered statistically significant.

#### Trajectory analysis and switch gene analysis

Pseudotime analysis was employed to infer the developmental trajectories of individual cells using transcriptomic data collected at different time points. This analysis provides valuable insights into cellular lineage trajectories and the role of specific genes in this process. For pseudotime analysis, we utilized the monocle2 and monocle3 packages ^47–49^ with default parameters. Separate analyses were performed on immune cells, melanocytes, and fibroblast-epithelial-peripheral cell populations.

Gene switch analysis is a method employed to identify genes that exhibit differential expression during cell differentiation or transformation, thereby potentially playing crucial roles in cellular development and differentiation. In this study, we utilized the GeneSwitches package ^50^ in R to identify transcription factors and cell surface proteins associated with gene switches, as well as their enriched pathways (GO, KEGG), using the results obtained from the pseudotime analysis of different developmental pathways. The findings obtained from this analysis provided valuable insights into the regulatory mechanisms underlying cell differentiation and highlighted potential targets for further investigation.

#### Cell communication analysis

Cell-to-cell communication is critical in various biological phenomena, including the pathogenesis of glaucoma. In this study, we utilized CellChat package ^51^ in R to investigate the communication between immune cells from patients with and without glaucoma. The computeCommunProb function was employed to identify ligand-receptor pairs involved in cell communication, while the computeCommunProbPathway function was used to summarize the communication patterns of these ligand-receptor pairs and their associated pathways. Moreover, to examine the differences in communication patterns between normal and glaucomatous samples, we integrated and compared the results using the mergeCellChat function. We also calculated the up- or down-regulated cellular communications across different sample types. Additionally, communication and pathway analyses were separately performed on fibroblasts, epithelial cells, and peripheral cells isolated from glaucoma patients, as the fibroblast-epithelial-peripheral cell populations primarily consisted of cells from patients with glaucoma.

## Results

### Results from TM

#### datasets Datasets Preprocessing

A total of 14,635 genes from 8 samples (normal n = 4, POAG n = 4) from GSE138125 , and 36 samples (normal n = 19, POAG n = 17) from GSE27276, were analyzed respectively. As shown in the PCA plot (**Suppl. Fig. 1C**), POAG and healthy control samples were separated into four distinct clusters (**Suppl. Fig. 1A**), indicating that there are differentially expressed genes among the four clusters.

#### GSEA for Potential Molecular Mechanisms

To gain a comprehensive understanding of pathologically enriched pathways of POAG within the hTM, we initially conducted GSEA analysis utilizing the GO and KEGG pathway databases (**Suppl. Fig. 2**). Enrichment maps were constructed to visualize the significantly enriched biological processes (BP) (**Fig. 2A**), molecular functions (MF) (**Fig. 2B**), cellular components (CC) (**Fig. 2C**), and KEGG pathways (**Fig. 3A**) that exhibited a strong association with POAG and interconnectedness with existing connections.

**Figure 2:**
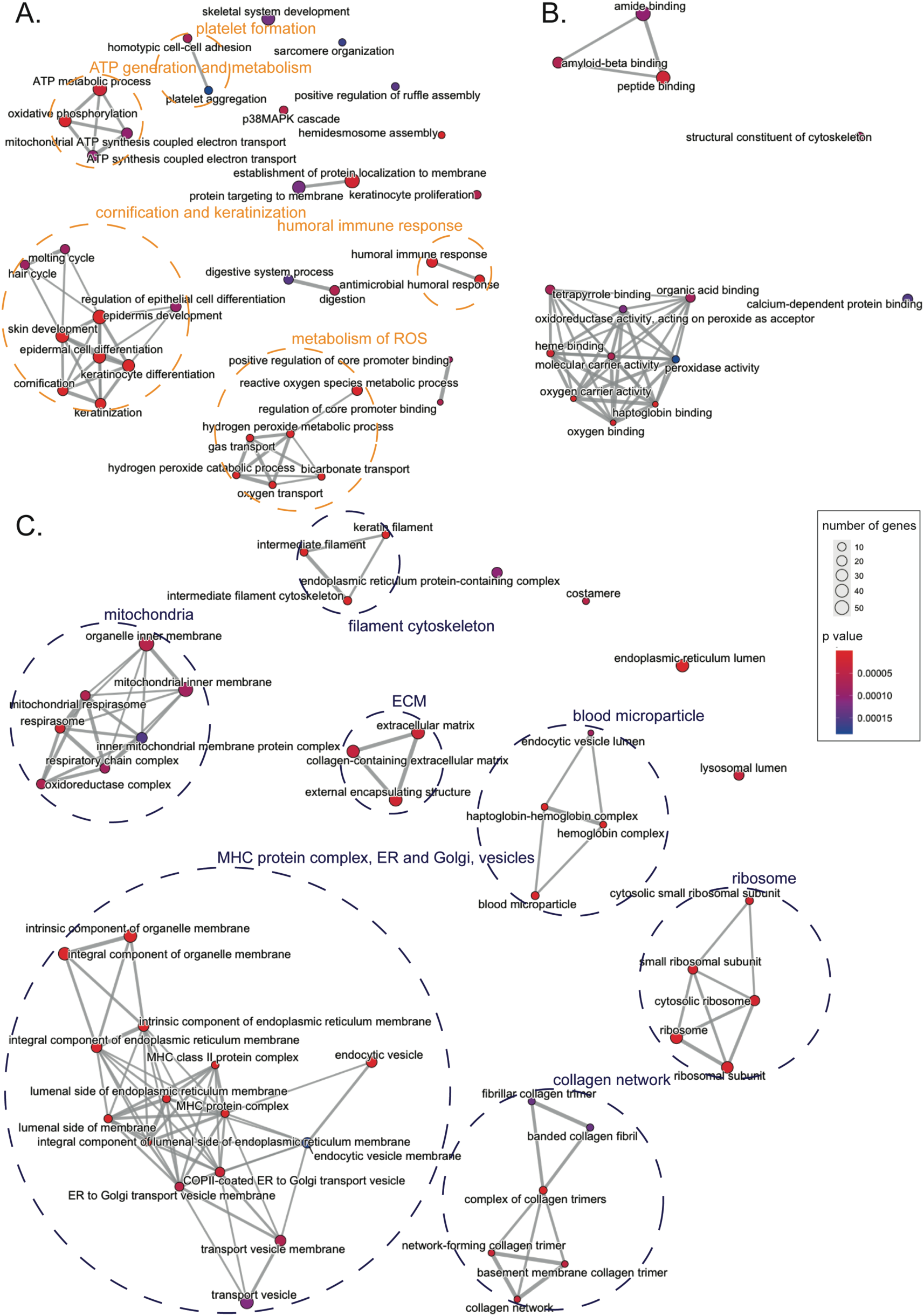
Gene Sets Enrichment Analysis (GSEA) Using Gene Ontology (GO) Database. **(A)**Enrichment map of GO annotations (BP: biological process). **(B)** Enrichment map of GO annotations (MF: Molecular function). **(C)** Enrichment map of GO annotations (CC: cellular component). Nodes represent enriched gene sets, grouped by similarity. Node size corresponds to the gene count in each set, and line thickness indicates shared genes between sets.

**Figure 3:**
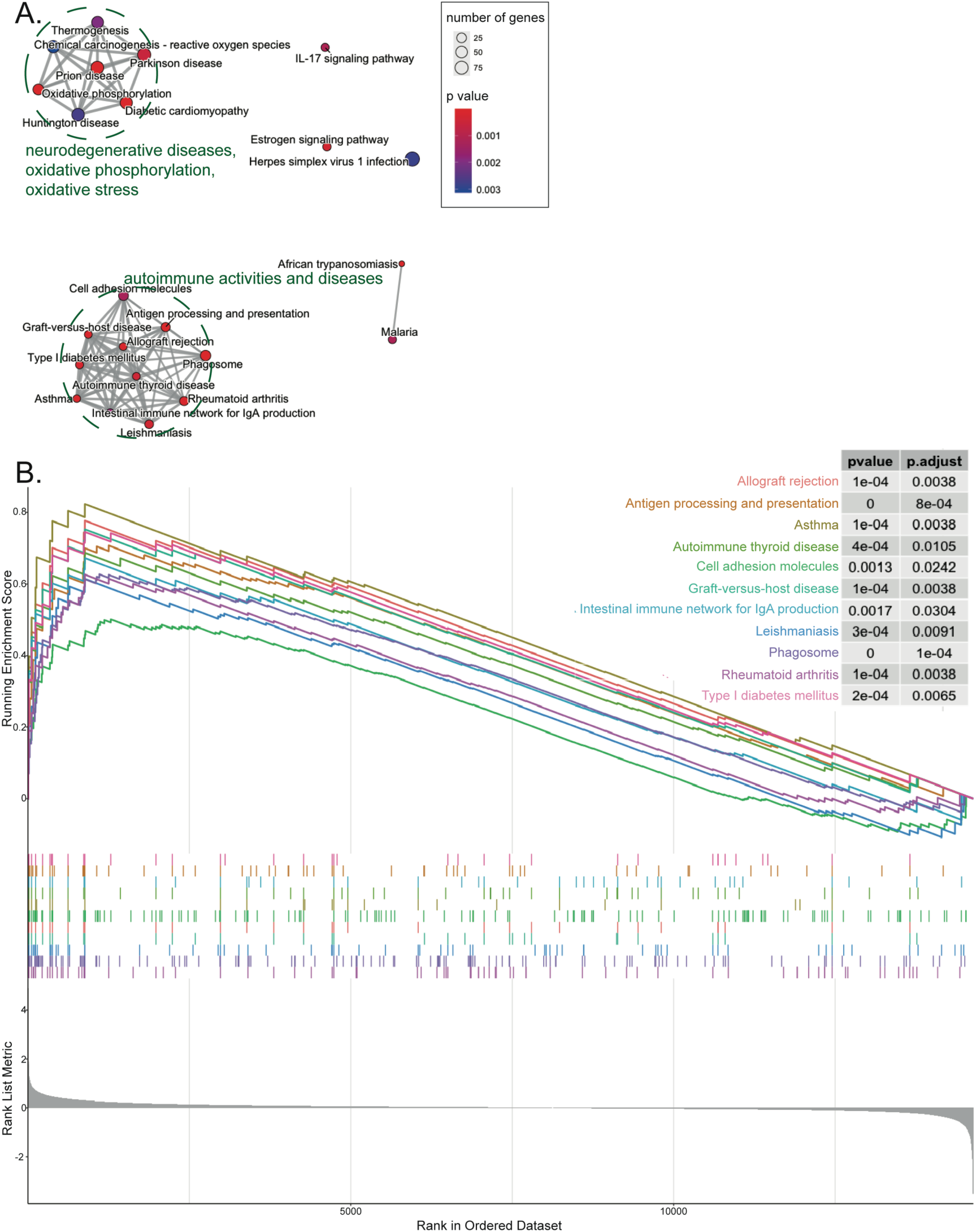
Gene Sets Enrichment Analysis (GSEA) Using Kyoto Encyclopedia of Genes and Genomes (KEGG) Database. **(A)**Enrichment map of KEGG pathways. **(B)** GSEA plot of immune-related KEGG pathways. In the enrichment map, nodes represent enriched gene sets grouped by similarity. Node size indicates gene set size, and line thickness represents shared genes between sets. In the GSEA plot, the top curves show running enrichment scores for each pathway, quantifying gene enrichment relative to a ranked list of all genes ordered by log2FC (bottom). Vertical lines denote the positions of pathway genes in the ranked list.

First and foremost, we identified five significantly enriched clusters utilizing the BP database. These clusters were associated with ATP generation and metabolism, cornification and keratinization, metabolism of reactive oxygen species (ROS), humoral immune response, and platelet aggregation. Subsequently, the enriched CC and MF terms revealed similar information. The annotated CC terms included components related to energy balance, ROS metabolism, and immune mediation such as mitochondria, vesicles, endoplasmic reticulum (ER) and Golgi complex, major histocompatibility complex (MHC) protein complex, ribosome, as well as ECM structures such as the collagen network and filament cytoskeleton. Furthermore, hemocyte fragments like blood microparticles and the haptoglobin-hemoglobin complex were also part of the enriched CC terms.

The enriched KEGG pathways primarily encompassed disease-related pathways, falling into two main categories: neurodegenerative diseases (e.g., Parkinson’s disease, Huntington’s disease) with associated alterations in oxidative phosphorylation and oxidative stress, and pathway implicated in multiple autoimmune diseases. As depicted in the GSEA plot of enriched immune-related KEGG pathways (**Fig. 3B**), these pathways were predominantly enriched in genes exhibiting relatively high expression levels in POAG.

#### Identification and functional analysis of DEGs

To conduct a more detailed examination of abnormalities, we identified 102 gene expression variations from the integrated dataset, 36 and 66 of which were upregulated and downregulated, respectively (**Fig. 4**). The heatmap of DEGs (**Fig. 4B**) demonstrated distinct classification of samples from the POAG and control groups although the PCA results (**Suppl. Fig.1 C**) indicated the existence of differences in the gene expression profiles between the two datasets.

**Figure 4:**
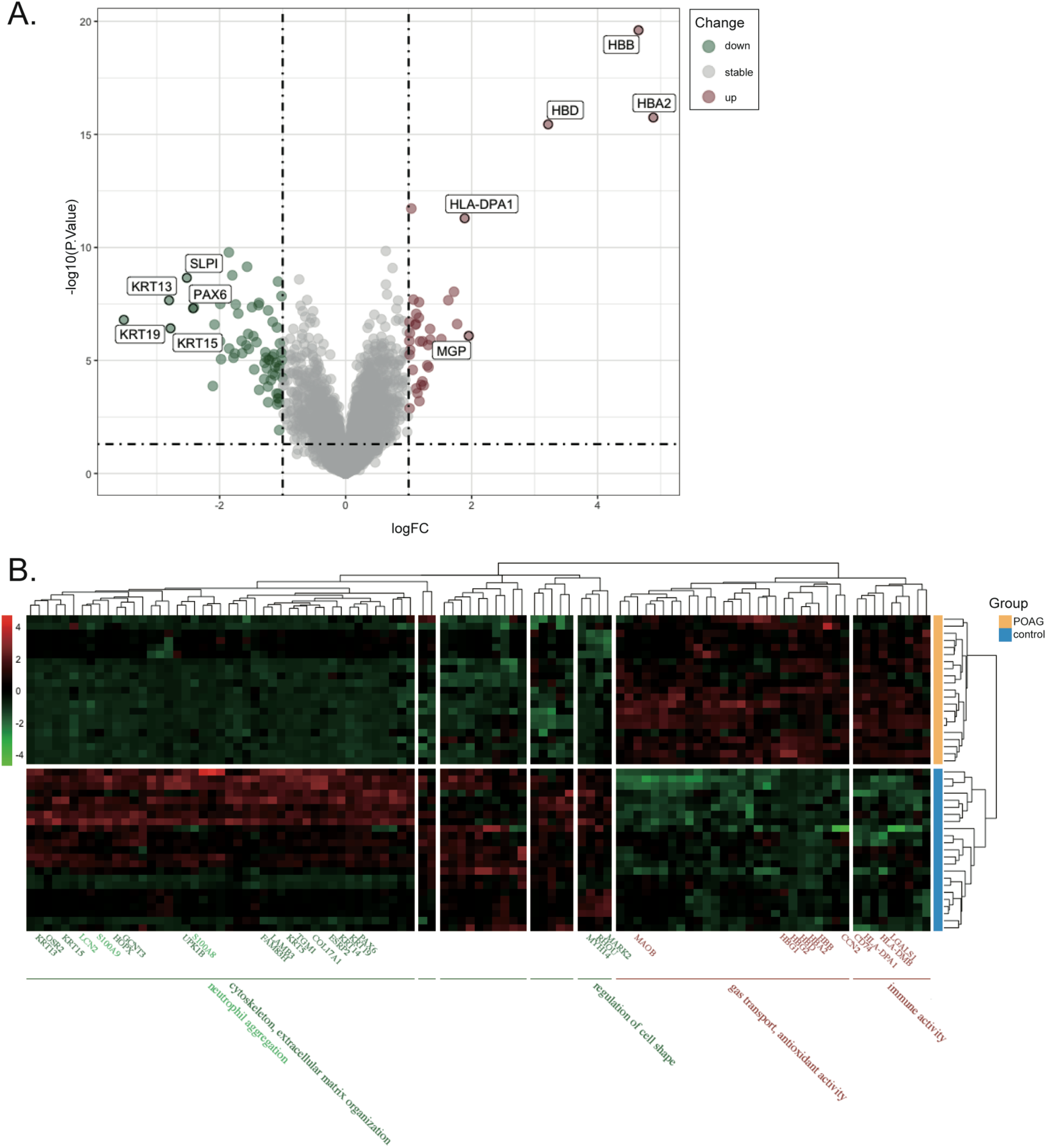
Differentially Expressed Genes (DEGs) in the Integrated Dataset. **(A)**Volcano plot of DEGs (cut-off values: *adj.P* < 0.05, |log2FC| > 1). **(B)** Heatmap of DEGs with GO-BP annotations, row-scaled at level 3. Red labels indicate upregulated genes, and green labels indicate downregulated genes in POAG.

To further investigate the biological implications of these DEGs, GO-BP annotations were performed for the row-scaled clusters at level 3. The analysis revealed that upregulated genes were associated with immune mediation, gas transport, and antioxidant activities. On the other hand, the downregulated genes were potentially involved in cell shape regulation, cytoskeleton and ECM organization, and neutrophil aggregation.

Furthermore, functional analyses encompassing GO-BP, CC, MF, and KEGG pathway functional analyses of all DEGs were conducted for all DEGs, yielding results consistent with the GSEA findings (**Fig. 3** and **5**). Notably, the majority of enriched KEGG pathways were related to autoimmunity, suggesting that changes in immune mediation play a significant role in POAG cases (**Fig. 5B**). For more detailed information, please refer to **Supplementary Tables 1** and **2**, which provide the results of GO enrichment and KEGG pathway analyses.

**Figure 5:**
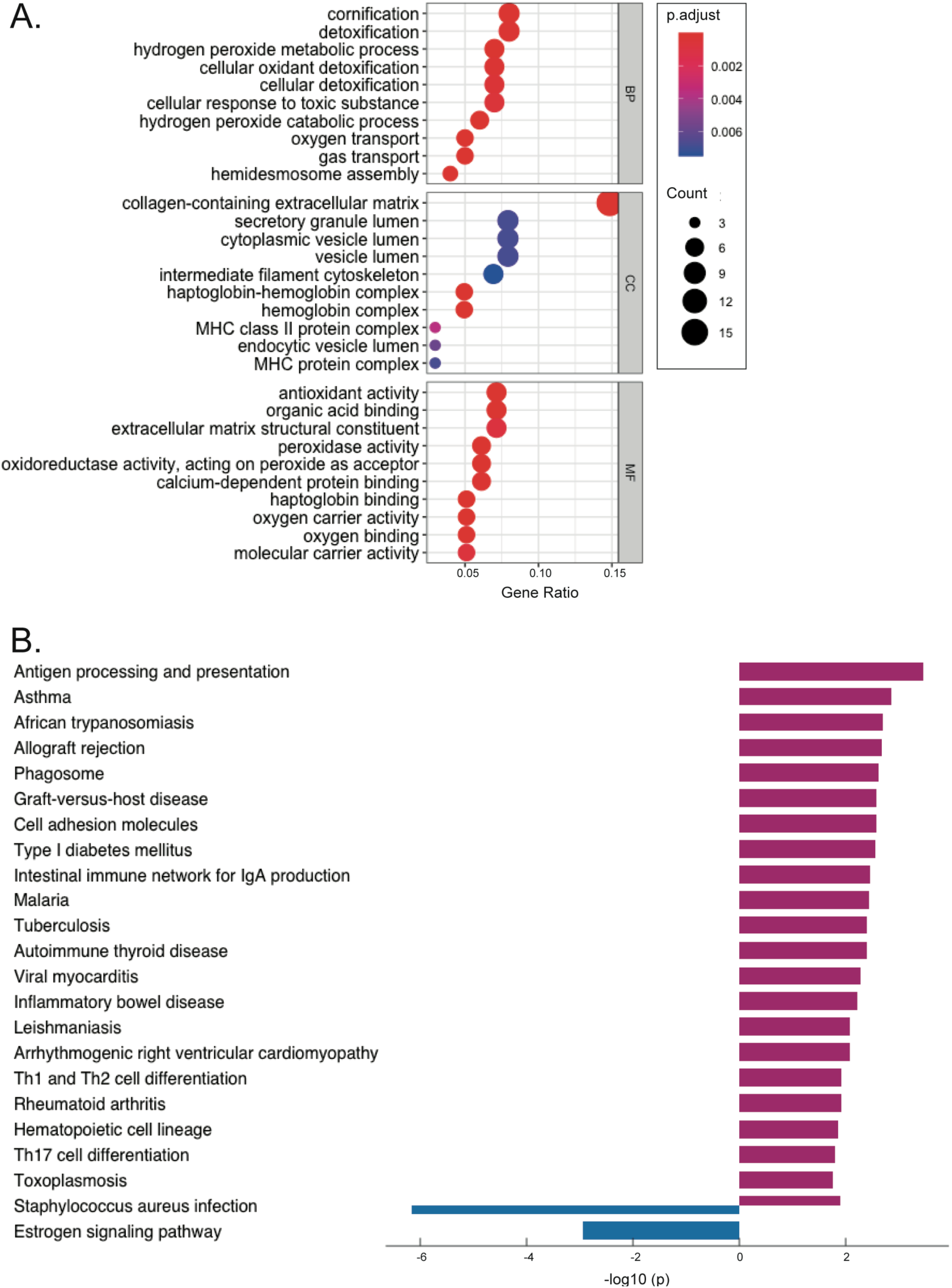
Enriched GO Annotations and KEGG Pathways of DEGs in POAG. **(A)**Top ten significantly enriched GO terms grouped into BP, CC, and MF functional categories. **(B)** Significantly enriched KEGG pathways.

#### PPI network and hub genes

To investigate the associations among DEGs in POAG, we uploaded all DEGs to the STRING database and constructed a PPI network (*P.Val* < 1.0^e-16^) comprising 102 nodes and 144 edges. The resulting network is presented in **Figure 6A**, where nodes with larger sizes and thicker lines indicate potentially key genes. In a PPI network, genes with more edges usually have a more critical role. Given the identification of approximately five categories of abnormalities through the GSEA and GO-KEGG pathway analyses of DEGs in the TM of patients with POAG, we selected the 30 most intersecting genes as potential hub genes to gain a relatively comprehensive understanding of the underlying pathology of POAG (**Fig. 6B**). The expression and function profiles of these hub genes are displayed in **Figure 7** and **Supplementary Table 3**.

**Figure 6:**
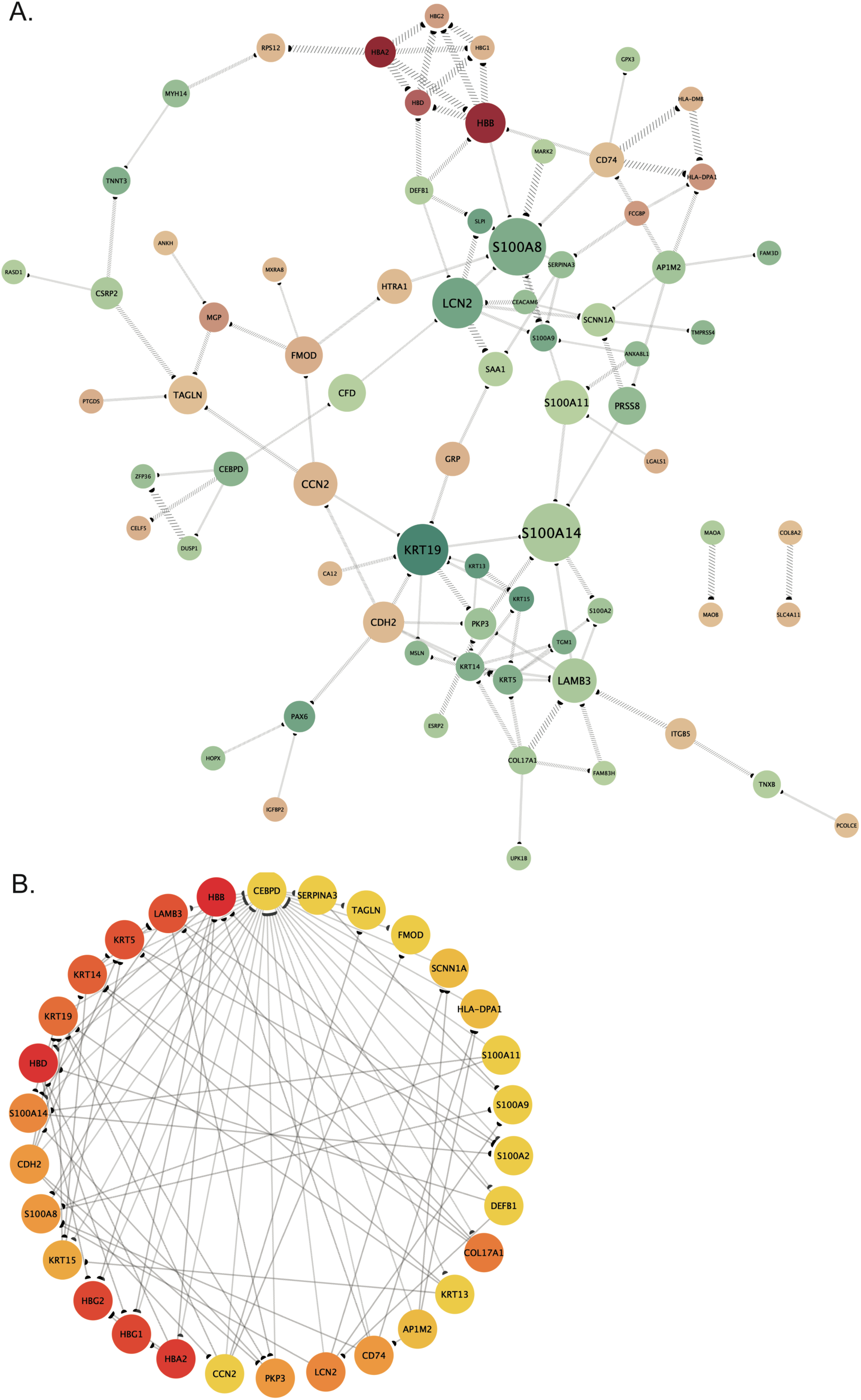
STRING and PPI Network of DEGs. **(A)**STRING interactions of DEGs, with node color representing log2FC, node size indicating cytoCNA betweenness, edge width reflecting cytoCNA combined score, and arrows indicating direction. **(B)** Hub genes: Top thirty genes ranked by MCC scores. Colors denote MCC score rank, and their arrangement is based on degrees (cytohubba), counterclockwise.

**Figure 7:**
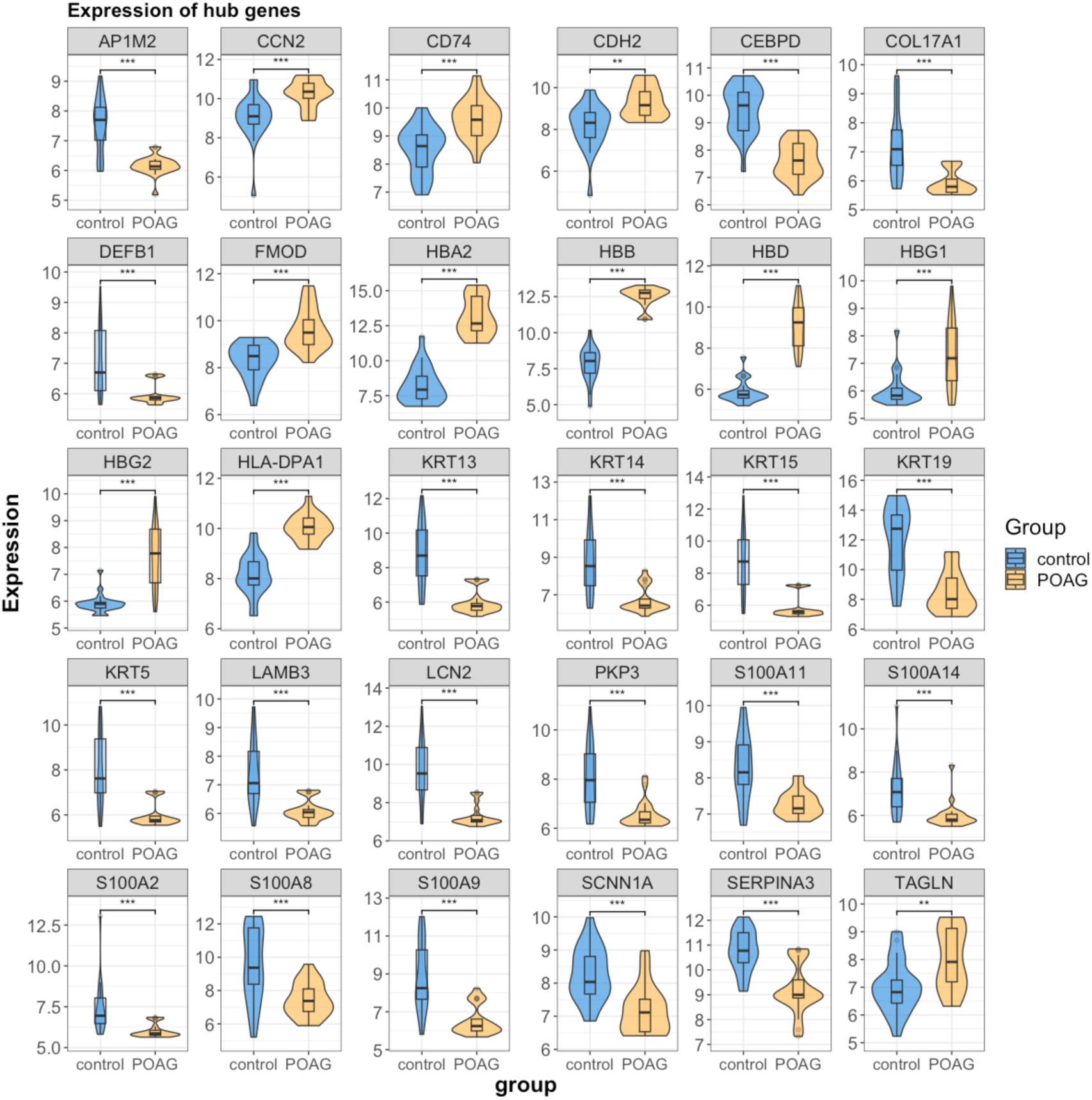
Expression Profiles of Hub Genes Among DEGs. Wilcoxon tests were conducted to assess the significance of expression differences.

To study the importance of these 30 potential hub genes, we conducted ROC analyses to evaluate their predictive efficiency in separating disease from control using gene expression level. As shown in **Figure 8**, all of these genes showed high accuracy on separating POAG cases from control groups, with area under the curve (AUC) values exceeding 0.75. This suggests their potential use as RNA biomarkers for POAG disease diagnosis. Notably, HBB (AUC = 1, 95% CI [1.000, 1.000]), HBD (AUC = 0.998, 95% CI [0.957, 1.000]), HBA2 (AUC = 0.992, 95% CI [0.957, 1.000]), and HLA-DPA1 (AUC = 0.979, 95% CI [0.913, 1.000]) displayed the highest AUC values. The first three genes are components of hemoglobin while the last one is a part of the MHC II complex, which plays a vital role in antigen presentation on the cell surface for recognition by CD4 T-cells ^52^.

**Figure 8:**
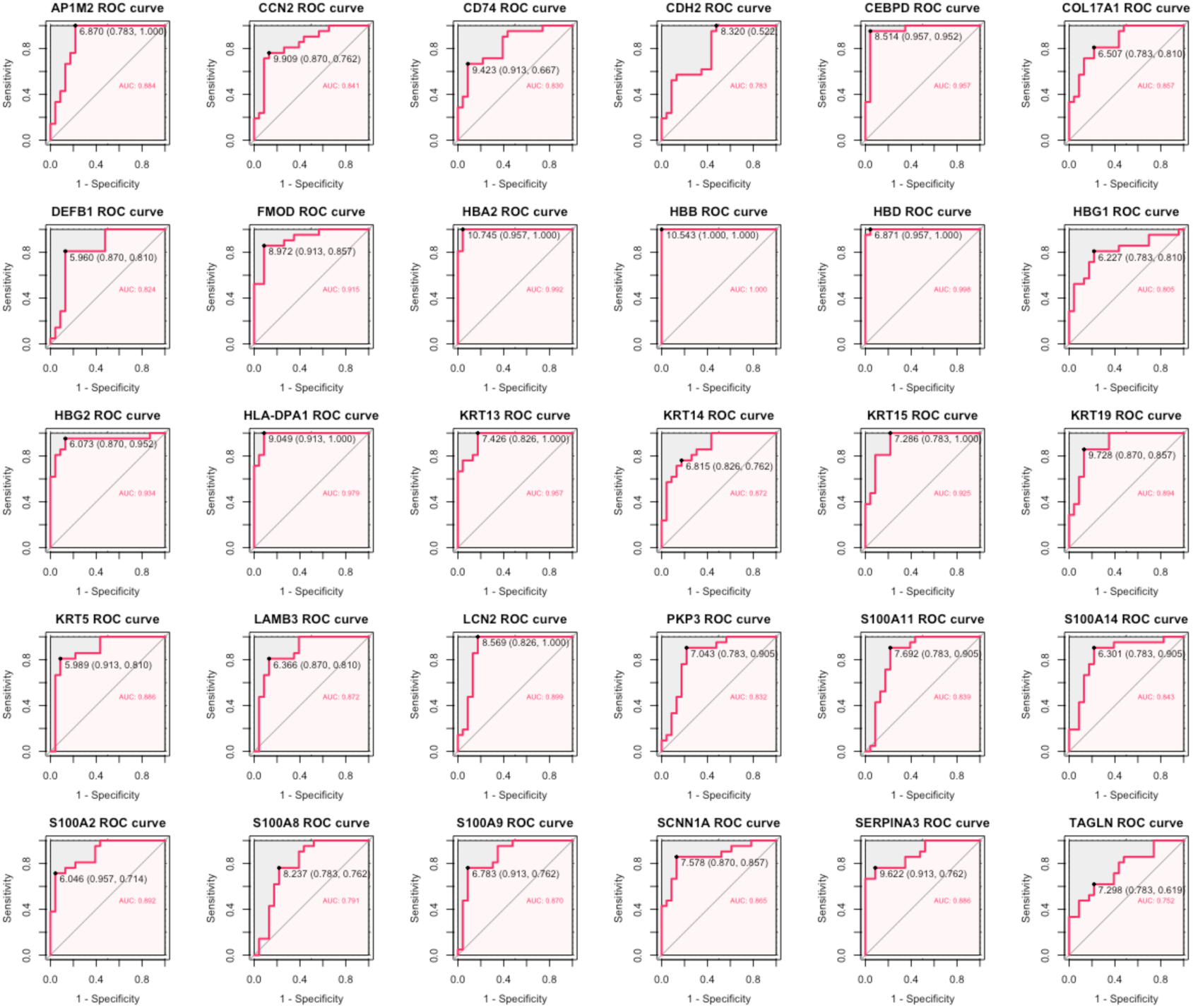
Diagnostic Accuracy of Hub Genes Among DEGs Assessed via ROC Curve Analysis.

#### Immune infiltration in POAG hTM

Next, MCP-counter analysis was employed to assess the differences among eight immune cell populations (T cells, CD8 T cells, NK cells, cytotoxic lymphocytes, B lineage, monocytic lineage, myeloid dendritic cells, and neutrophils) and two stromal cell populations (endothelial cells and fibroblasts) in POAG and normal samples (**Fig. 9**). Interesting, results revealed a higher proportion of T cells, but not CD8^+^ T cells, NK cells, monocytic lineage, endothelial cells, and fibroblasts in POAG compared with normal hTM (all *P.Val* < 0.05) (**Fig. 9B**). This suggests a certain level of immune activation and tissue hyperplasia in POAG. Furthermore, the increased proportion of monocytic lineage cells, but not myeloid dendritic cells, in POAG samples indicates the infiltration of macrophages. It is worth noting that although not statistically significant, there was a tendency towards decreased neutrophil infiltration in the POAG group, which aligns with the findings obtained from the GO-BP annotations of row-scaled clusters of the down-regulated DEGs (**Fig. 4B**).

**Figure 9:**
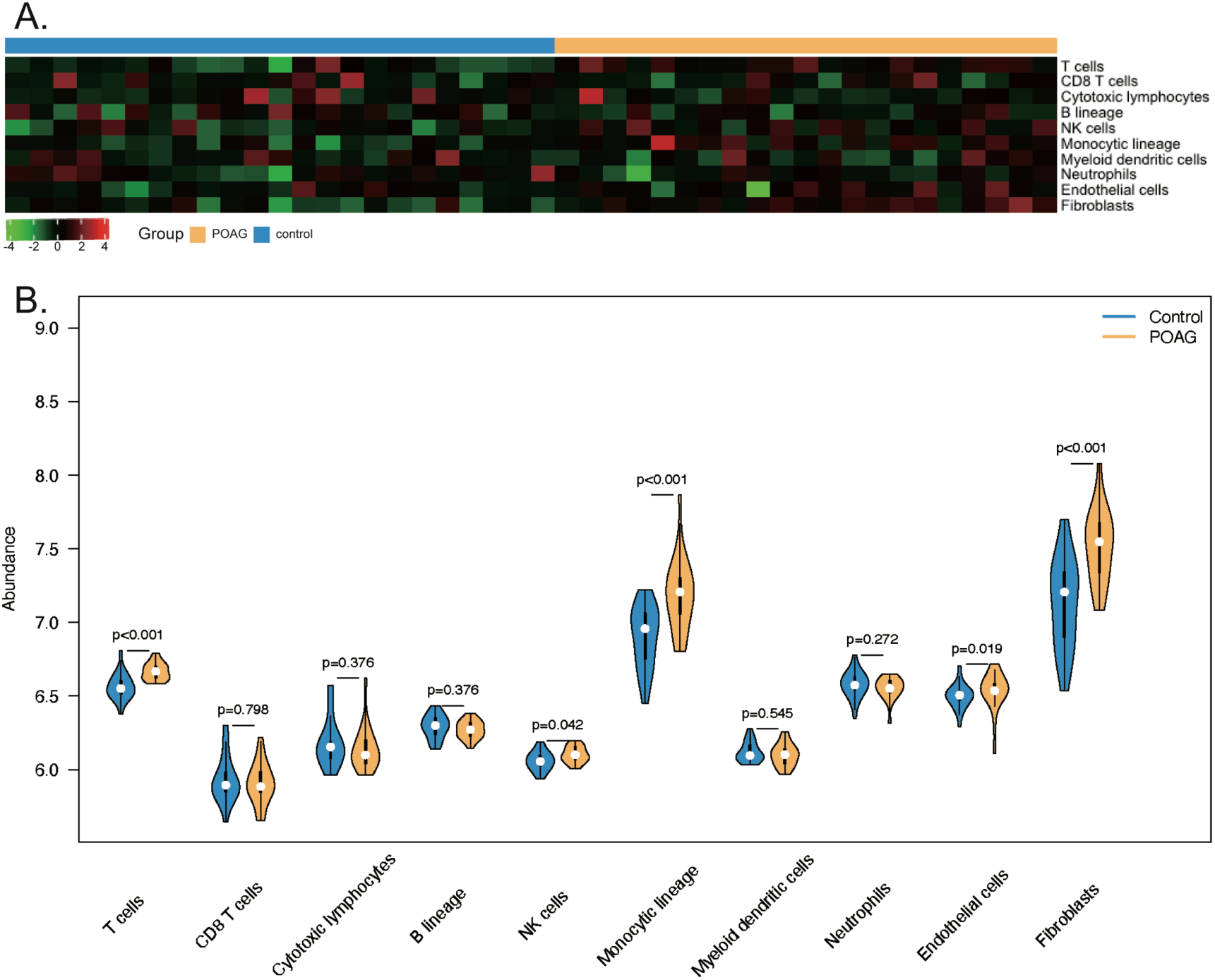
Exploration of the Immune Landscape using the Integrated Dataset. **(A)**Heatmap displaying immune cell types. **(B)** Comparison of immune cell type fractions between control and POAG groups, with significance assessed through Wilcoxon tests.

#### Identification of POAG-associated WGCNA modules and their relationship with immune infiltration

To identify genes closely associated with POAG, we performed WGCNA and set the soft threshold to 4 to ensure the network exhibited a scale-free characteristic. An outlier control sample (GSM674419) from the GSE27276 dataset was removed, and the remaining samples were used to construct the weighted network (**Fig. 10AB**). Subsequently, average linkage hierarchical clustering based on topological overlap matrix (TOM) differences and dynamic tree pruning yielded nineteen modules, denoted by various colors (**Fig. 10C**). Among the five robust POAG-related WGCNA modules (|Cor| > 0.5, *P.Val* < 0.05), the midnightblue module (77 genes; r = 0.62, *P.Val* < 0.001) and the lightcyan module (50 genes; r = 0.64, *P.Val* < 0.001) exhibited the strongest positive and negative correlations with POAG, respectively (**Fig. 10D**).

**Figure 10:**
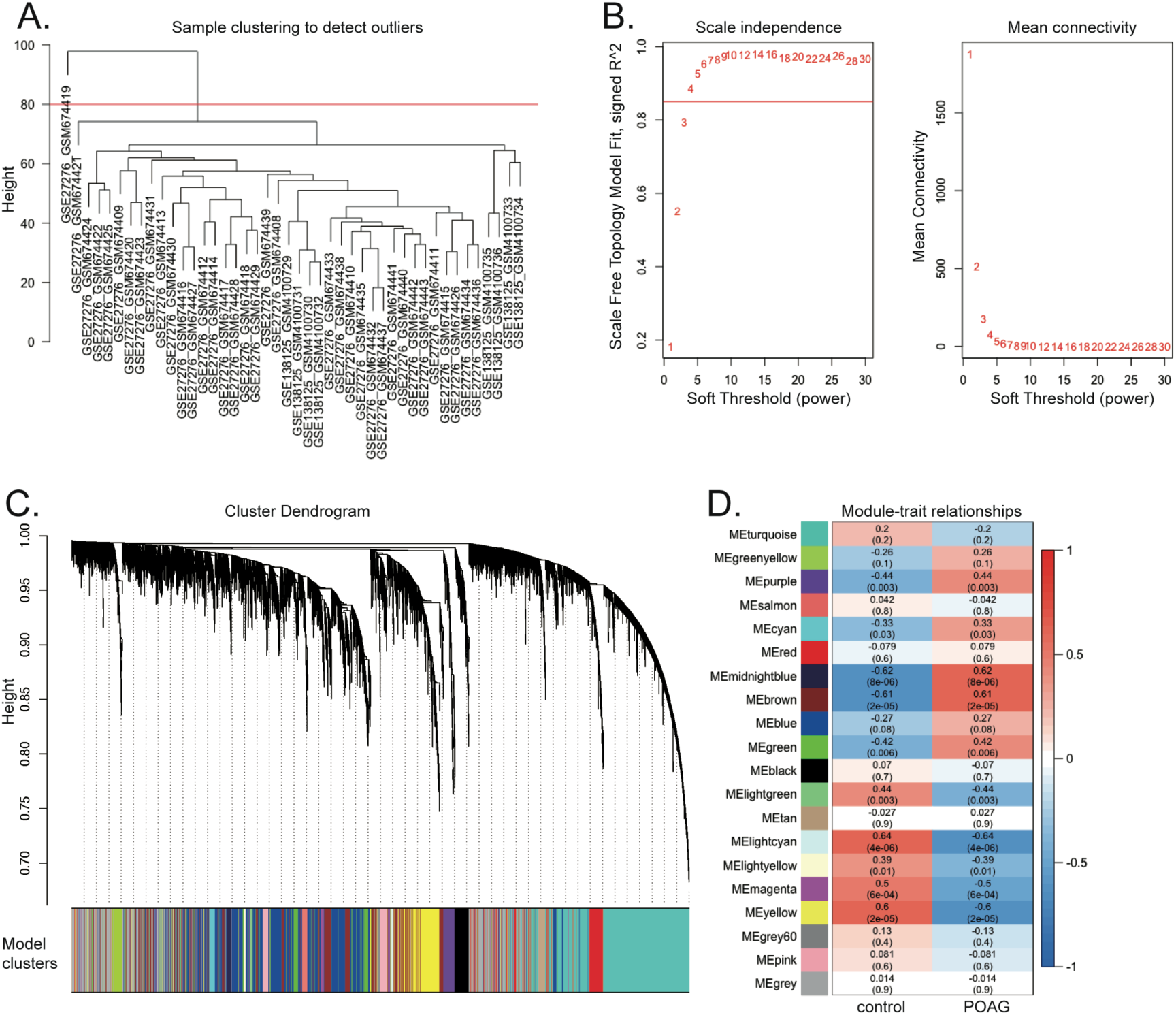
Weighted Gene Co-Expression Network Analysis (WGCNA) using the Integrated Dataset. **(A)**Sample clustering to identify outliers. **(B)** Scale-free fitting index of soft threshold power and mean connectivity. **(C)** Gene dendrogram obtained through average linkage hierarchical clustering, with each color representing a gene module. **(D)** Correlation between gene modules and POAG, with numbers indicating correlation coefficients (p-values) on the horizontal and vertical axes.

Next, we assessed the correlation between all identified WGCNA modules and the immune cell scores obtained from the MCP-counter analysis (**Fig. 11**). Interestingly, both the top POAG-related WGCNA modules, midnightblue (r = 0.35, *P.Val* = 0.02) and lightcyan (r = -0.59, *P.Val* < 0.001), showed the closest relationship with T-cell infiltrations. Additionally, they demonstrated statistically significant associations with monocytic lineage and fibroblasts infiltrations (all *P.Val* < 0.05). Module eigengene-based connectivity was then calculated to identify hub genes within these two modules, considering their absolute kME value higher than 0.8. Subsequently, the 14 hub genes from the midnightblue module (**Fig. 12A**) and the 10 hub genes from the lightcyan module (**Fig. 13A**) were uploaded to the STRING database to construct PPI network. The overlapping genes primarily involved immune regulation (**Suppl. Table 4** and **5**). GO annotations for the hub genes of the positively POAG-related module revealed enrichment in biological processes related to the positive regulation of lymphocyte proliferation and chemokine production. The cellular components included the MHC II complex, components of endocytic vesicles, ER, and Golgi apparatus, while the molecular function annotation was associated with MHC II protein complex binding (**Fig. 12B**). Conversely, hub genes from the negatively POAG-related module were mainly involved in the inactivation of MAPK activity (**Fig. 13B**). Therefore, it can be inferred that genes from both the top POAG-related WGCNA modules potentially contribute to immune activation.

**Figure 11:**
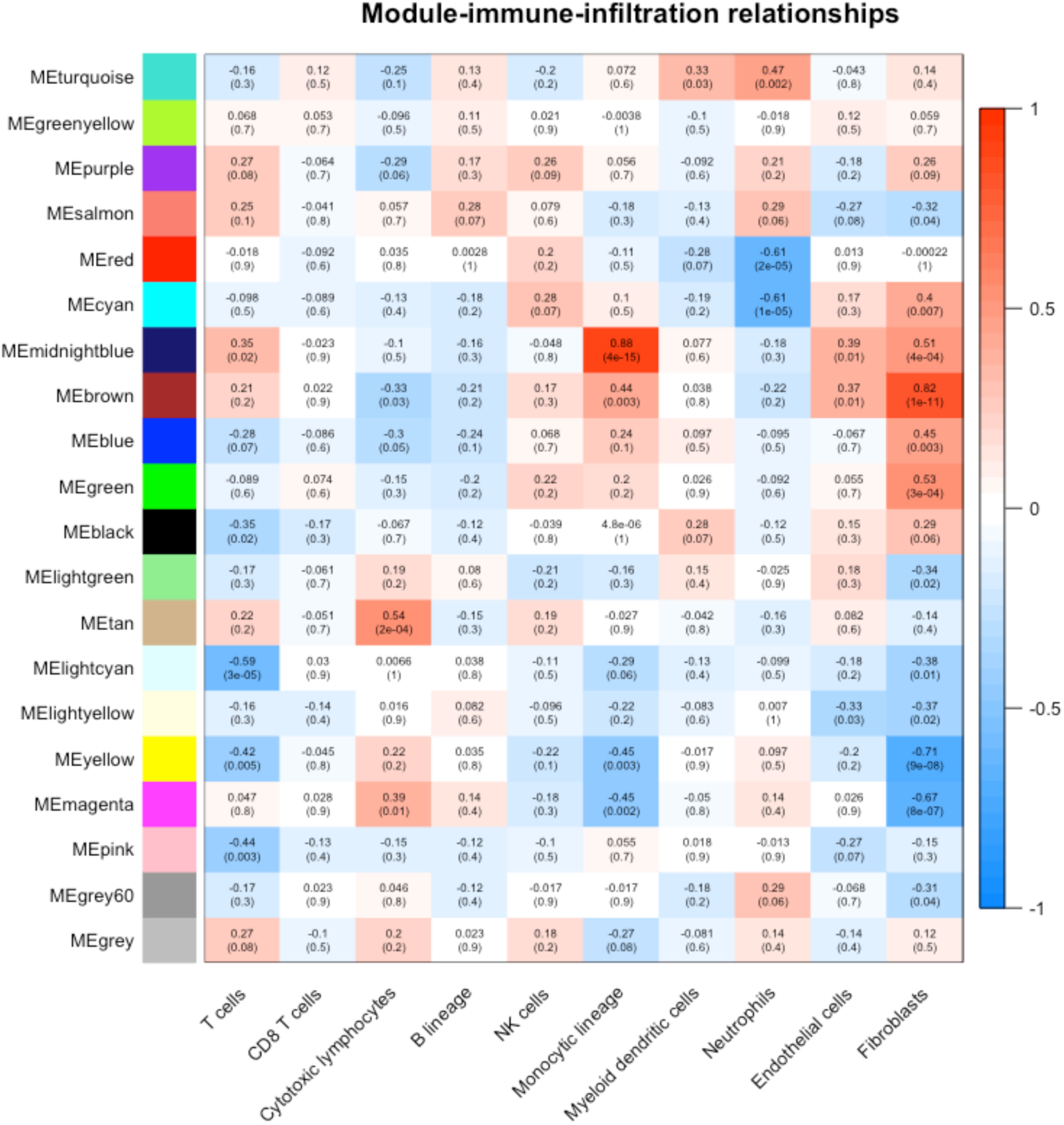
Correlation between Gene Modules and POAG-Related Immune Infiltrates. Numbers represent correlation coefficients (p-values) on the horizontal and vertical axes, indicating the strength and significance of the correlation between gene modules and immune infiltrates in the context of POAG.

**Figure 12:**
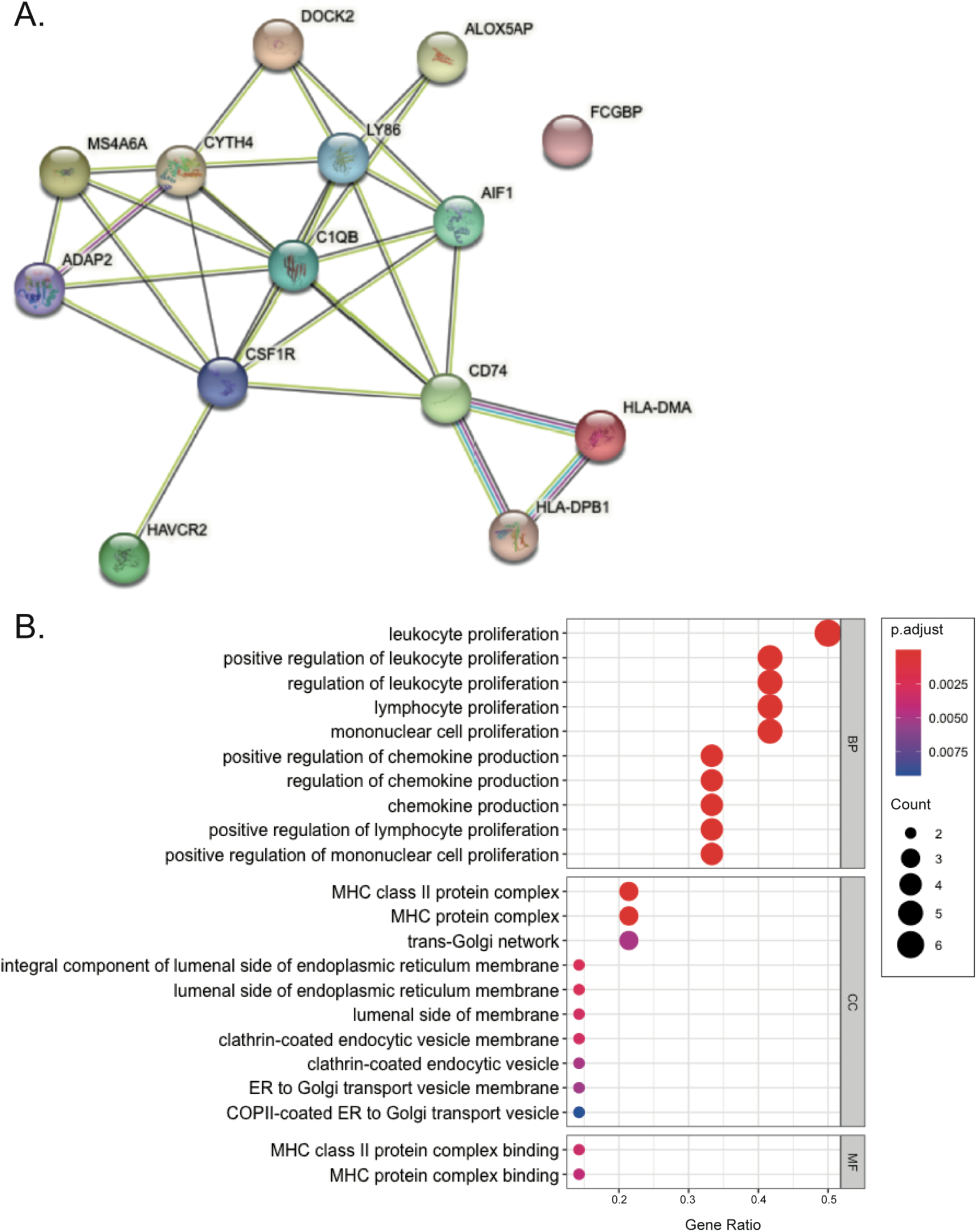
STRING Interactions and GO Annotations for Hub Genes in Positively POAG-Related Modules. **(A)**STRING interactions of hub genes from the “midnightblue” module. **(B)** GO annotations of hub genes from the “midnightblue” module.

**Figure 13:**
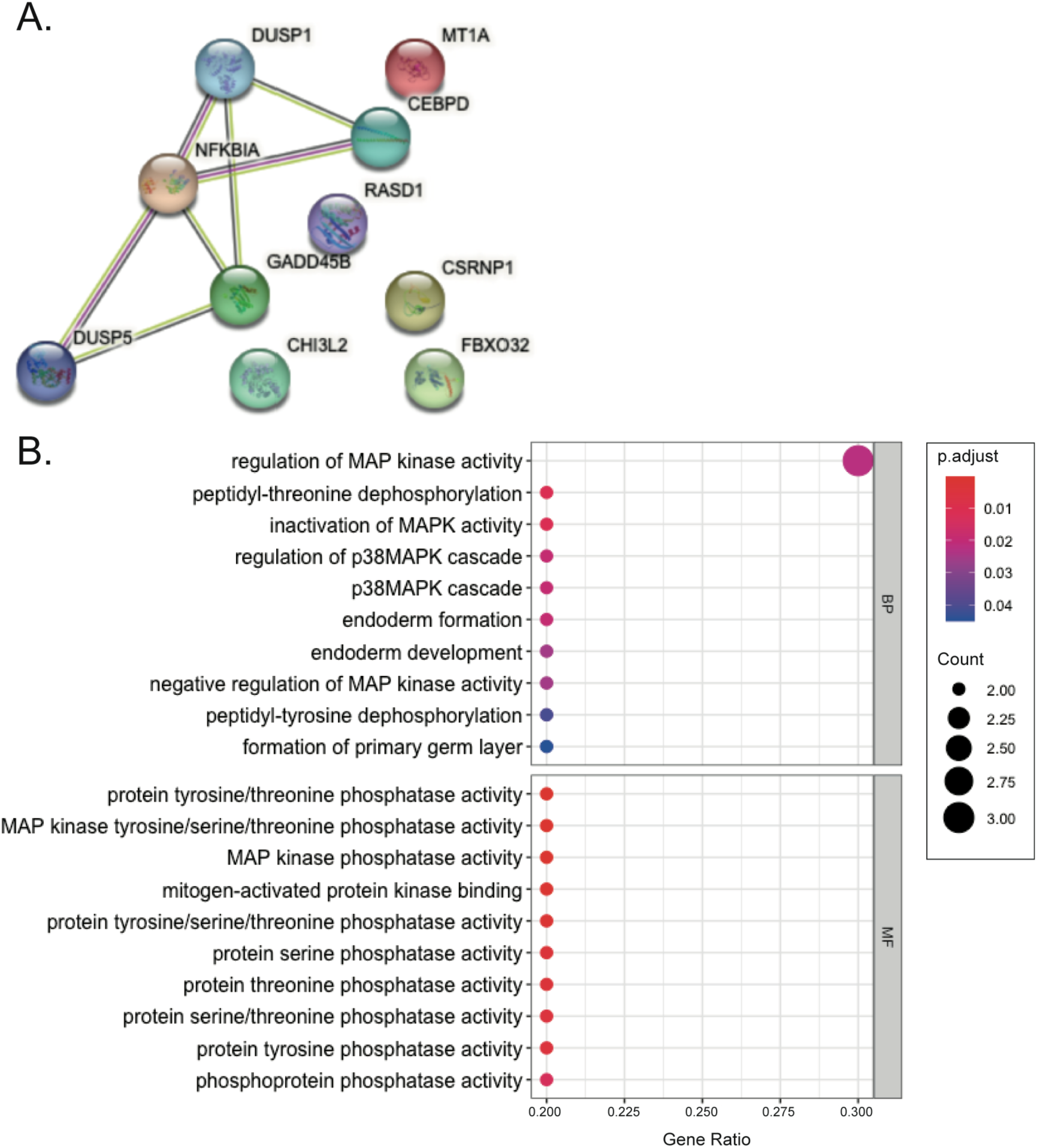
STRING Interactions and GO Annotations for Hub Genes in Negatively POAG-Related Modules. **(A)**STRING interactions of hub genes from the “lightcyan” module. **(B)** GO annotations of hub genes from the “lightcyan” module.

### Single-cell RNA-sequencing data analysis using choroid dataset

#### Cell atlas of global glaucoma cells in the choroid

We conducted a single-cell RNA-sequencing data analysis of choroidal tissues from patients with glaucoma to investigate the expression profiles of different cells. Our study included disease samples from glaucoma and AMD, whereas controls consisted of individuals with early-stage AMD only. Using marker genes provided by the cellmarker2.0 database ^47^, we classified all cells in the global single-cell view of glaucoma into 14 distinct cell types (**Fig. 14F**). These cell types encompassed five immune cell types: B cells, leukocytes, macrophages, mast cells, and natural killer T cells (NKT cells); and three types of epithelial cells: choriocapillaris, vein endothelial cells, artery endothelial cells; two types of Schwann cells, two types of peripheral cells, fibroblastic cells, and melanocytes, respectively (**Fig. 14A**). Each cell type exhibited unique expression patterns specific to its respective cell cluster (**Fig. 14D**).

**Figure 14:**
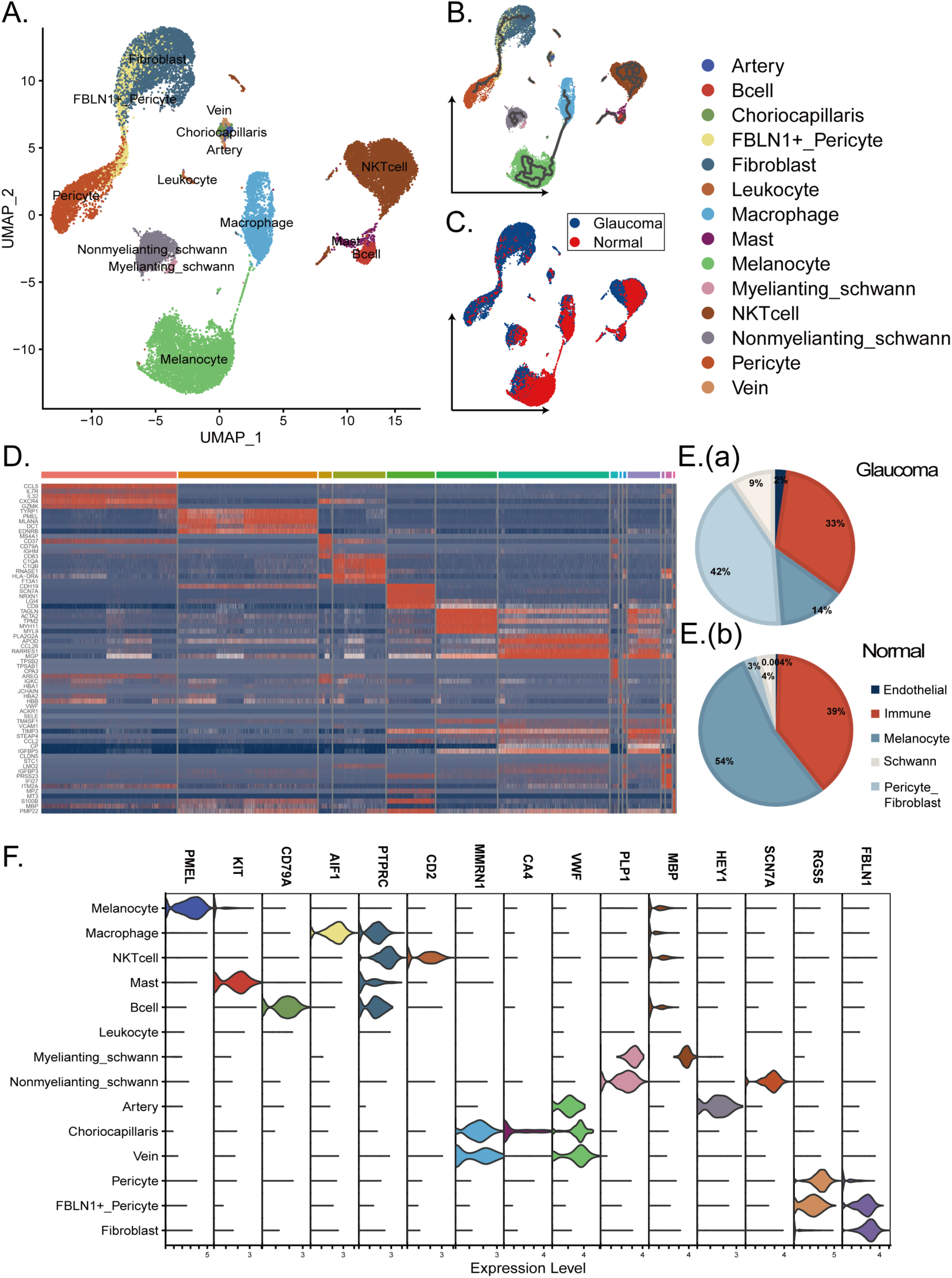
Single-Cell Analysis of Glaucoma and AMD Patients. **(A)**The cell annotation results of single cells mapping glaucoma by UMAP including melanocyte, immune cells (macrophage, NKT cell, B cell, leukocyte and mast cell), schwann cells (nonmyelianting schwann cells and myelinating schwann cells), pericyte cells (pericyte and FBLN1+ pericyte), fibroblast, endothelial cells (choriocapillaris, artery endothelial cells and vein endothelial cells). **(B)** The UMAP visualization with cell pseudotime reconstruction by monocle3. **(C)** The UMAP visualization of different diagnosis type, including glaucoma group (patients who have glaucoma and early AMD) and normal group (patients who only have early AMD). **(D)** Heatmap of marker genes in different cell types. **(E)** Pie charts of the cell proportion of glaucoma group (a) and normal group (b), in which pericyte fibroblast mixed up as one type. **(F)** Volcano plot of marker genes from CellMarker2.0 in different cell types.

During our analysis of the descending results, we made an intriguing observation of a previously unidentified cell type positioned between fibroblasts and peripheral cells. This particular cell type exhibited high expression levels of both fibroblast marker (FBLN1) and peripheral cell marker (RGS5) (**Fig. 14F**). Recognizing the potential implications of pericyte-fibroblast transformation (PFT) on the inflammatory response and disease-related enrichment pathways ^53^, we redefined this cell type as FBLN1^+^ peripheral cells. Our analysis indicated that these cells represented transitional state cells undergoing PFT with some degree of proliferation in glaucoma choroid tissue (**Fig. 14B**).

Furthermore, we conducted a comparison of the cellular composition between control and glaucomatous patients (**Fig. 14C**). We observed that the largest fraction of cells in patients with glaucoma originated from peripheral cells, fibroblasts, and FBLN1^+^ pericytes, while melanocytes were the most abundant in the control group. The analysis of cell proportion revealed significant differences in cellular composition between the control and glaucoma groups (**Fig. 14E**).

In conclusion, our single-cell profiling of the choroid tissues in glaucoma patients has provided valuable insights into the variations in cellular components within the microenvironment. These findings enhance our understanding of glaucoma’s pathophysiology and have the potential to drive the development of novel diagnostic and therapeutic strategies in the future.

#### Trajectory analysis and identification of switch genes for melanocytes

Loss of vision is a typical symptom in glaucoma, and melanocytes, which are light-sensitive cells in the human eye, may contribute to vision loss ^54^. In our study, we observed a significantly lower percentage of melanocytes relative to the total cell population in glaucoma patients (0.137) compared to control individuals (0.538). This finding aligns with previous observations of reduced light sensitivity in glaucoma patients. To unravel the molecular mechanisms underlying melanocyte-related vision loss in glaucoma, we focused on the melanocyte cell clusters and performed trajectory analysis ^48^. Our analyses revealed a trajectory from normal cells to glaucoma cells, with a concentration of cells in the pseudotime fraction greater than 8 (**Fig. 15AB**).

Moreover, we identified 45 switch genes, including 35 differentially expressed genes (logFC > 0.58), 5 transcription factors (NR2F1, DDIT3, NME2, KLF2, MAFB), and 2 cell surface proteins (TNFRSF14, TYRP1) (**Fig. 15C** and **E**). Additionally, we found significant enrichment of NF-kB, MAPK, and protein folding processes (**Fig. 15D**), which have been previously associated with melanin synthesis in melanocytes ^55–57^. However, their relationship with glaucoma in melanocytes remains unclear. We postulate that the downregulation of these pathways in glaucoma patients may lead to decreased melanin synthesis, resulting in reduced light sensitivity of melanocytes and subsequent vision loss, including symptoms of blindness. Furthermore, we identified two pathways related to the heat response (GO:0009408 and GO:0034605) (**Fig. 15D**), which have not been previously reported in the context of glaucomatous melanocyte lesions. These findings suggest that these biological processes may be involved in the pathogenesis of glaucoma and warrant further investigation.

**Figure 15:**
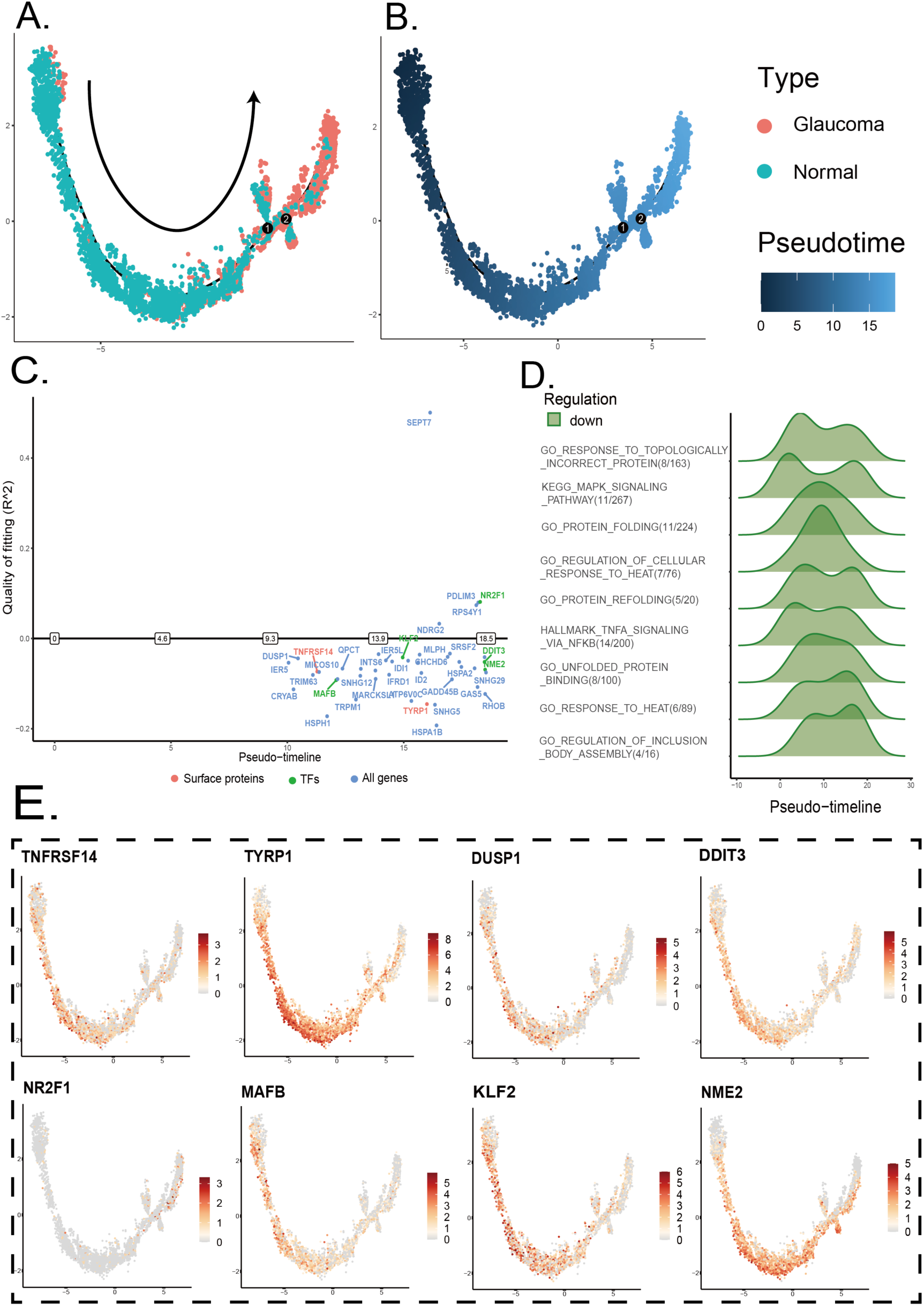
The Pseudotime Result and Switch Gene Detection Result of Melanocyte in Choroid. **(A)**The cell trajectories of melanocyte analyzed by monocle2, labeled by the diagnosis result of samples. **(B)** The cell trajectories labeled by the pseudotime analyzed by monocle2. **(C)** The switch genes with pseudotime greater than 8 in the process of transferring from normal to glaucoma melanocytes. **(D)** The cell enrichment abundance analysis based on the switch genes in the melanocyte. **(E)** The expression level of switch genes in melanocyte, including 5 transcription factors (NR2F1, DDIT3, NME2, KLF2, MAFB), and 2 cell surface proteins (TNFRSF14, TYRP1) and DUSP1.

#### ECM-related pathways in pericyte fibroblast transfer

Choroidal thickening has been commonly observed in glaucoma patients, and choroidal thickness (CT) is often considered an essential indicator of choroidal health ^58^. However, the molecular mechanisms underlying this process at both the single-cell and genetic levels remain unclear. To address this gap, we conducted temporal sequence and cellular communication analyses focusing on peripheral, fibrotic, and endothelial cells. Our analysis revealed an evolutionary pathway consistent with pericyte-fibroblast transition (PFT) (**Fig. 16AB**), characterized by high expression levels of both FBLN1 and RGS5 marker genes. In our study, we defined these cells as FBLN1^+^ fibroblasts (**Fig. 14A** and **16AB**).

**Figure 16:**
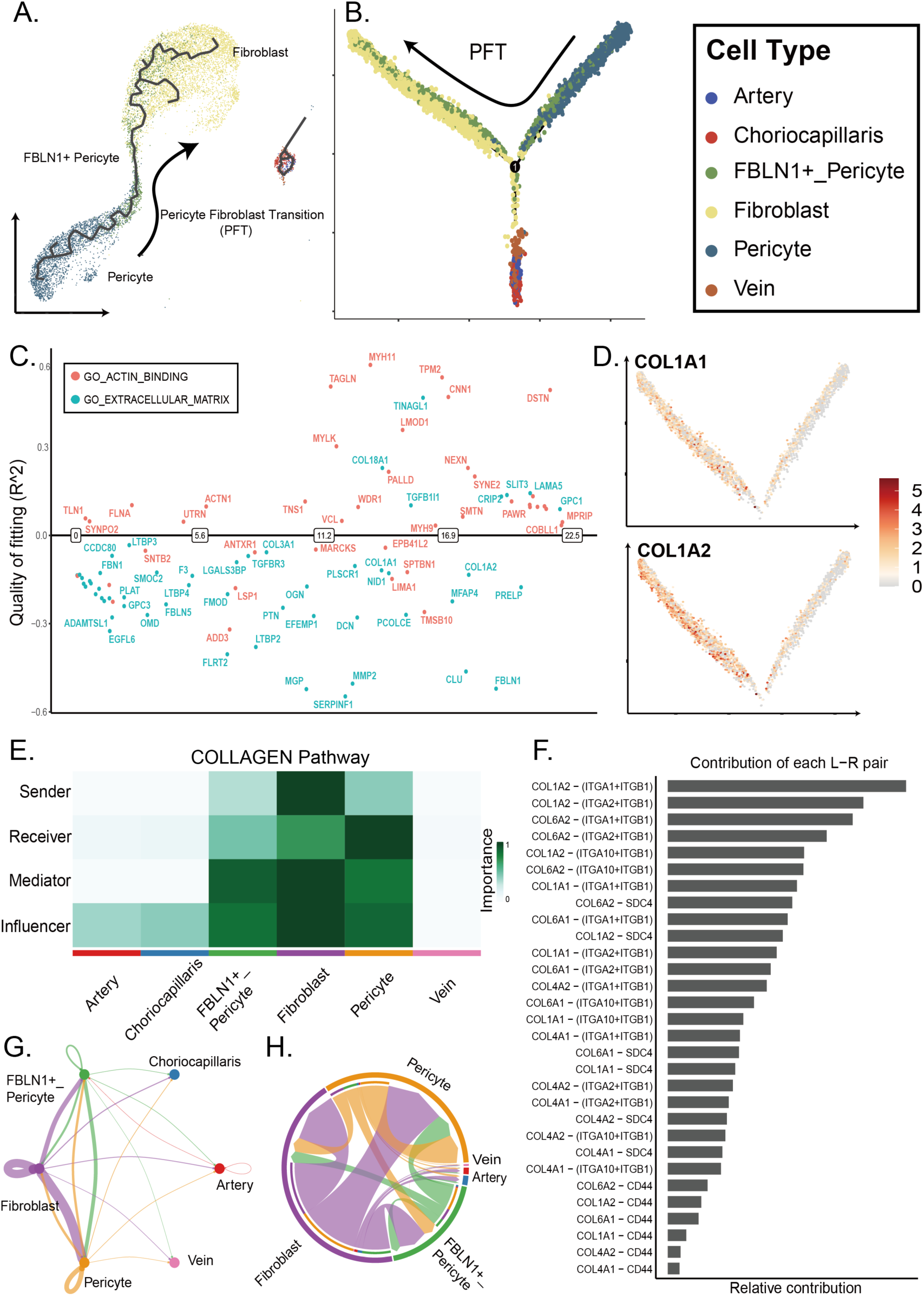
Analysis of Pericyte Fibroblast Transfer and Cell Communication in the Collagen Pathway. **(A)**The cell trajectories of pericyte fibroblast transfer in UMAP visualization based on monocle3. **(B)** The pseudotime analysis of PFT and endothelial cells based on monocle2. **(C)** The dot plot of switch genes in the extracellular matrix pathway and actin binding pathway in the PFT development process. **(D)** The cell expression abundance of COL1A1 and COL1A2 in the PFT process, COL1A1 and COL1A2 are the surface proteins in switch genes. **(E)** The heatmap with the role of different cell types in the cell communication of collagen related pathway base on CellChat. **(F)** The contribution of each ligand-receptor pair in the communication of collagen pathway. **(G, H)** The circle plot and net plot of the strength of cell-to-cell communication in collagen related pathway.

In this work, we identified 105 differentially expressed genes (logFC > 0.58), including 6 transcription factors and 24 cell surface proteins. Enrichment analysis of these genes revealed a concentration on extracellular matrix (ECM) and actin-binding pathways (**Fig. 16C**). Furthermore, the switch genes and surface proteins intersecting with the ECM pathway were COL1A1 and COL1A2 (**Fig. 16D**), which encode collagen type I, an important component of the ECM ^59^. To further explore and validate the role of the ECM pathways, we analyzed cell communication related to collagen type I in the collagen pathway. Our analysis showed that fibroblasts acted as ligands, peripheral cells as receptors, and FBLN1^+^ peripheral cells as intermediate transmitters in the communication network, which coincided with the development of PFT (**Fig. 16E-H**). Therefore, our findings suggest that the increased expression of COL1A1 and COL1A2 in fibroblasts may modify the physical properties of the ECM (**Fig. 16F**) by synthesizing collagen I, thereby influencing the PFT process and leading to choroidal thickening.

These results provide novel insights into the molecular mechanisms underlying choroidal thickening in glaucoma patients and have the potential to contribute to the development of new diagnostic and therapeutic strategies for this condition.

#### Intersection genes between the TM and choroid datasets

To investigate the relationship between changes in the anterior and posterior IOP-controlling eye segments with glaucoma, we combined the findings from the TM and choroid datasets. As shown in **Table 1**, 111 changed genes overlapped between the TM and choroid datasets and were referred to as intersection genes. These genes were either DEGs in the TM dataset or module genes related to POAG. Furthermore, they may be DEGs (markers) in any cell cluster (immune cells, endothelial-fibroblast-pericytes, and melanocytes) in the choroid single-cell dataset, switch genes of cell types, or ligands or receptors of cell communication. The number of plus signs (TRUE count) assigned to each gene in the table indicates the gene’s potential importance in the glaucoma context. To gain further insights into these highly significant genes, we utilized gene-integrated information from reputable databases such as NCBI (https://www.ncbi.nlm.nih.gov/ gene/) and GeneCard (https://www.genecards.org/).

**Table 1.**
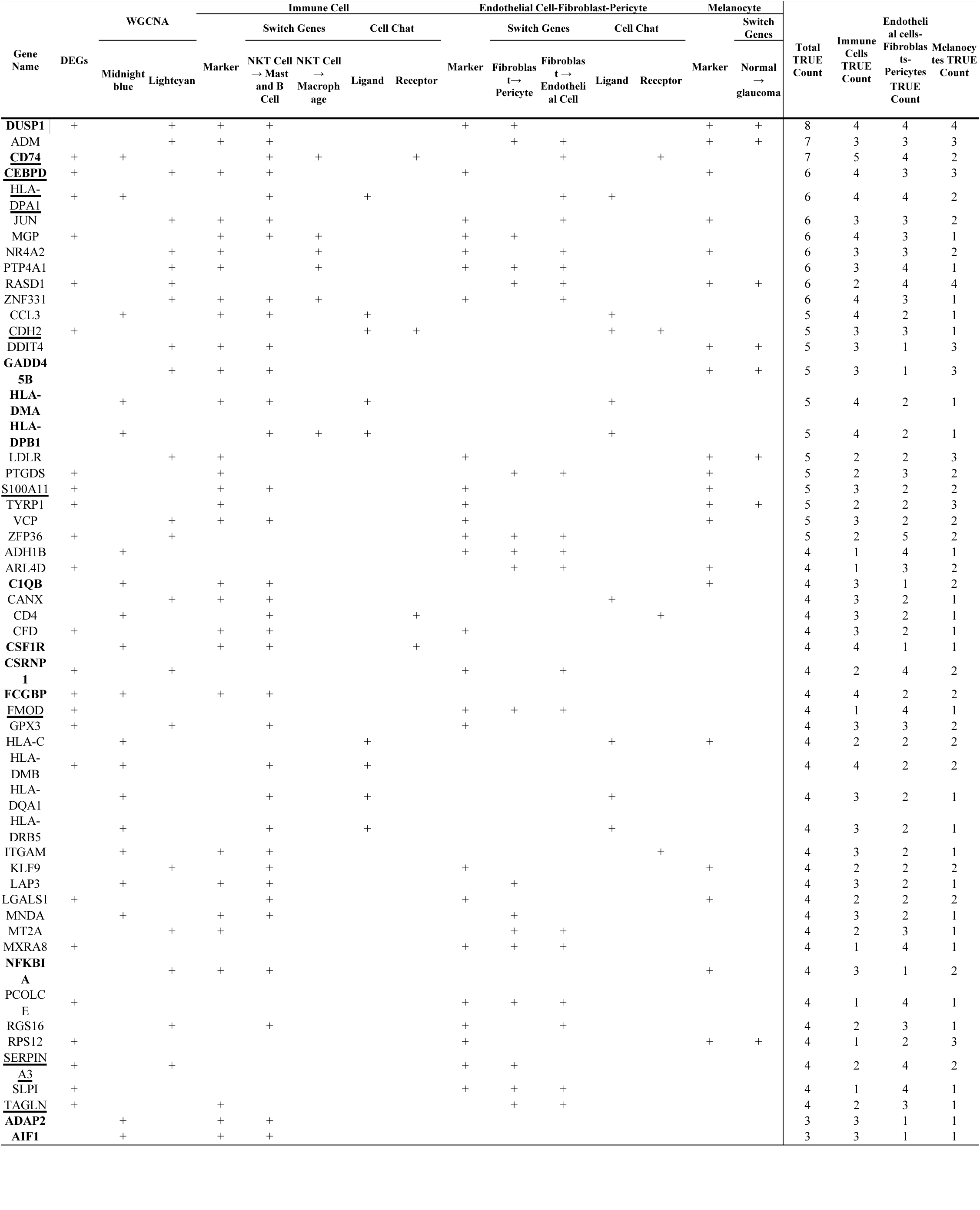
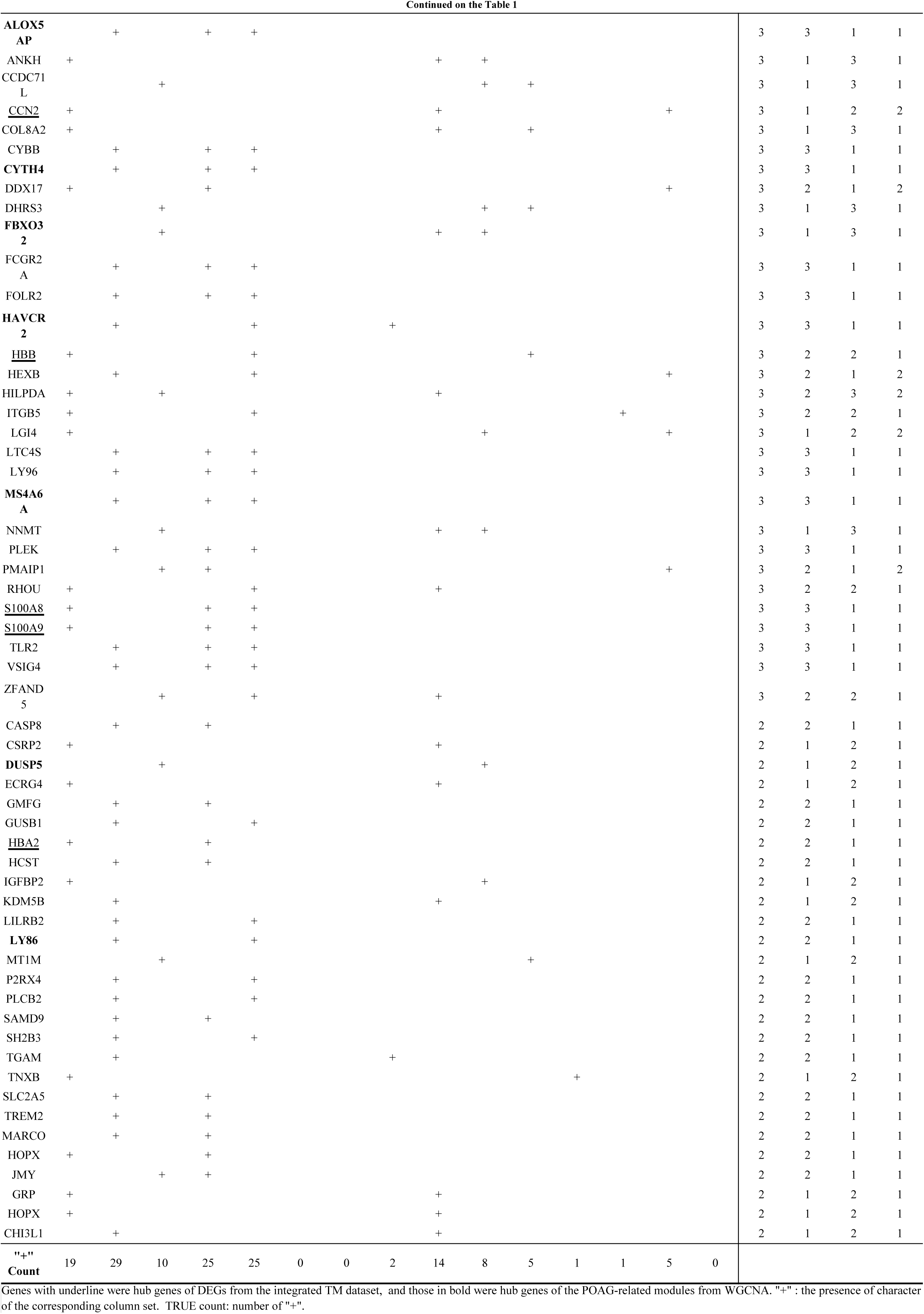
Gene intersection of TM and choroid datasets.

Among the identified genes, DUSP1 exhibited the highest TRUE count and served as a marker for all three major cell clusters. It also emerged as a hub gene in the negatively associated WGCNA module specific to POAG in the TM dataset. Known for tis role as a negative regulator of the MAPK pathway, DUSP1 acts as a phosphatase for tyrosine and threonine residules ^60,61^. The second-highest TRUE count was attributed to ADM, which has been implicated in hemodynamics and may have antibacterial properties ^62^. It is a switch gene in all three major cell clusters, indicating the transition from a normal to glaucoma state in melanocytes. In immune cells, it may have influenced the impact of NKT cell on mast and B cells, while in the endothelial-fibroblast-pericyte cluster, it may have played a role in the transition from fibroblast to pericyte (refer to **Table 1**).

Other genes with a TRUE count equal to or greater than 6 included CD74, CEBPD, HLA-DPA1, JUN, MGP, NR4A2, PTP4A1, RASD1, and ZNF331. According to information from NCBI and GeneCard, many of these genes are involved in immune and inflammatory responses, with some being implicated as biomarkers for neurodegenerative and autoimmune diseases. For example, JUN can induce inflammation, while CD74, CEBPD, HLA-DPA1, and ZNF331 are associated with immune and inflammatory responses. NR4A2 has been recognized as a biomarker for neurodegenerative and autoimmune disorders. PTP4A1, MGP, and RASD1, respectively, play roles in maintaining plasma membrane integrity, regulating vasculature calcification, and influencing ECM interactions, and cell morphology.

#### Enhancement of cell communication represented by the MHCII pathway

In the immune cell cluster, the intersection genes with high TRUE counts were mainly the same as those mentioned above with high total TRUE counts (refer to **Table 1**), further emphasizing the significant impact of immune response changes in glaucoma pathogenesis. Notably, the majority of genes with the highest TRUE counts were constituents of MHCII, including CD74, HLA-DPA1, HLA-DMA, HLA-DPB1, and HLA-DMB, indicating substantial alterations in the MHCII pathway within the immune cell cluster under glaucoma conditions (refer to **Fig. 17 A-B**). The HLA-D family acts as ligands in this pathway, whereas CD4, which is also an intersecting gene, is the receptor. As shown in **Figure 17 C-D**, the expression levels of both the HLA-D family members and CD4 increased.

**Figure 17:**
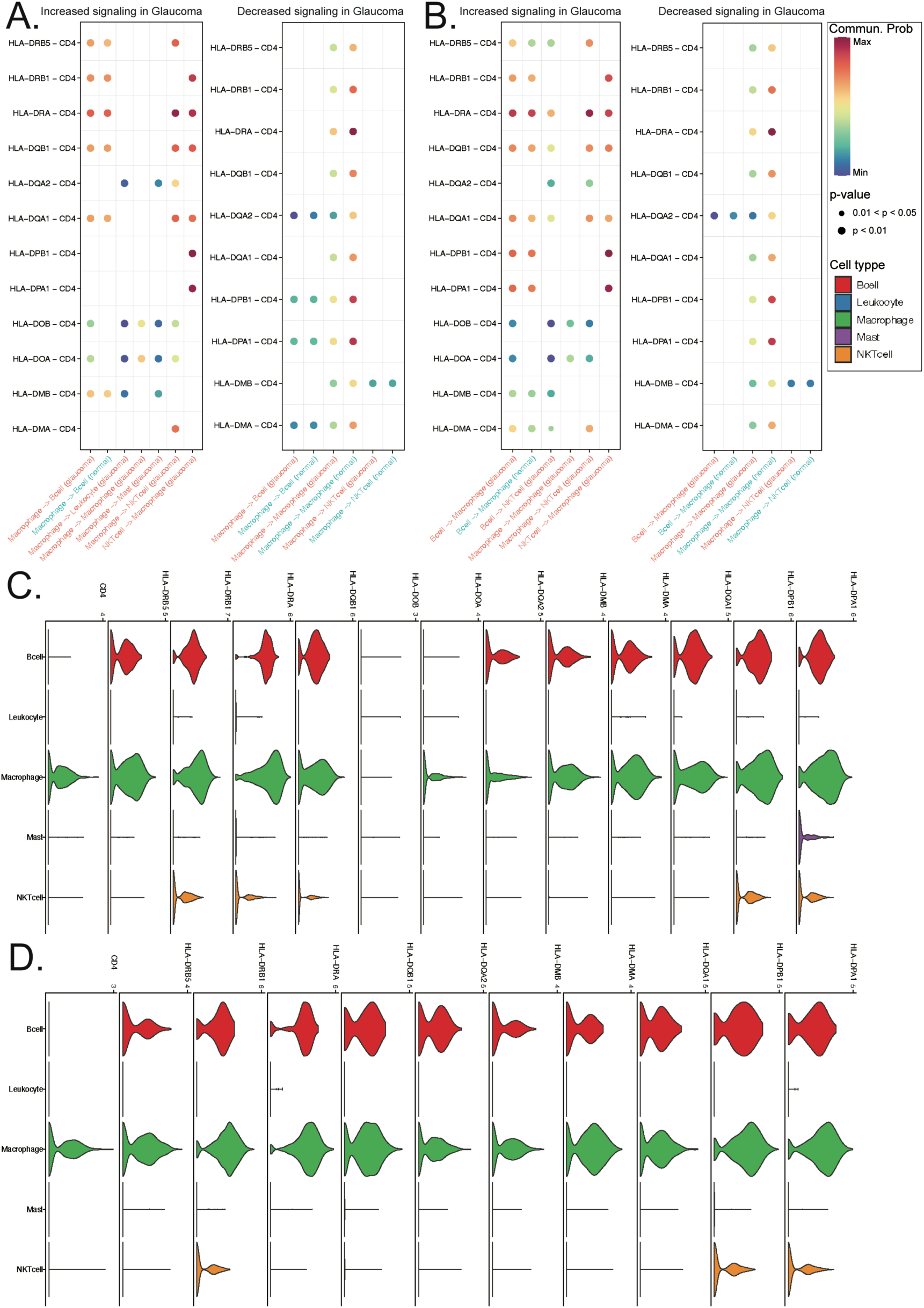
Comparison of Cell Chat in the MHCII Signaling Pathway Network Between Glaucoma and Normal Samples. **(A, B)** Differential cell communication regulated by the ligand and receptor, respectively. **(C, D)** Expression profiles of the cell chat genes in glaucoma and normal samples, respectively.

Pseudotime analysis of the immune cell cluster of the choroid data revealed trajectories from NKT cells to mast and B cells, as well as to macrophage (see **Figure 18 A-B**). Consistently, there was enhanced cell communication among immune cells, particularly the impact of NKT cells, with their role as influencers being the most significant (refer to **Figure 18 C-F**). Apart from their involvement in the MHCII signaling pathway network, several intersection genes served as ligands or receptors for intercellular communication, contributing to complex changes in communication among immune cells. These genes included CDH2, CCL3, HLA-C, CD74, CSF1R, CD4, ITGAM, HAVCR, and others (refer to **Table 1**). These genes may affect cell-cell contact or secreted signaling pathways such as CDH, MIF-CD74-CXCR4, CCL, MHCI, CSF, IL16, CD40LG-ITGAM-ITGB2, C3-ITGAM-ITGB2 (complement), GALECTIN, APP, FCER2-ITGAM-ITGB2 (CD23), and ICAM1-ITGAM-TIGB2 (ICAM).

**Figure 18:**
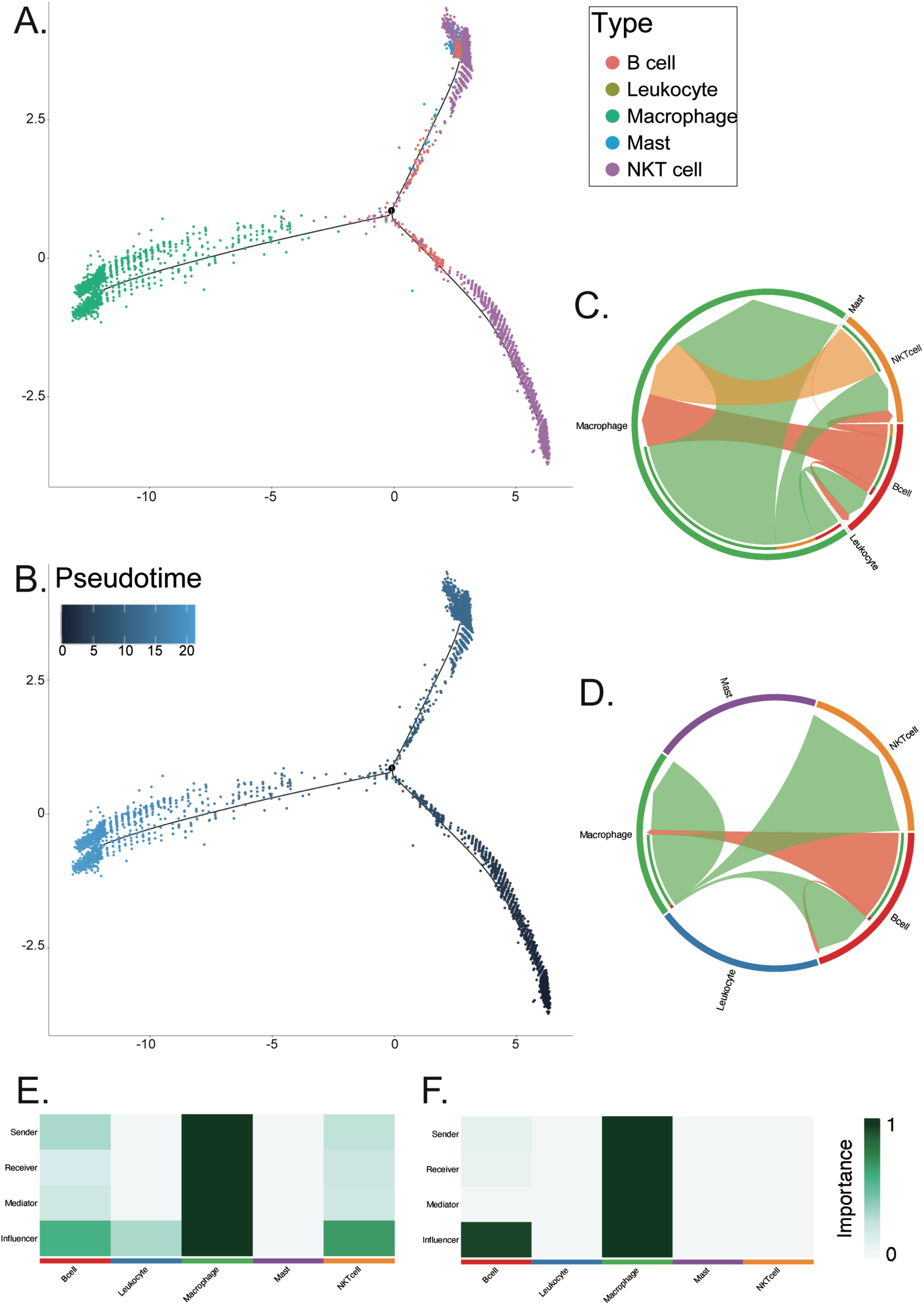
Analysis of Immune Cells and Cell Communication in the MHCII Signaling Pathway Network. **(A)**The cell trajectories of immune cells analyzed by monocle2, labeled by the diagnosis result of samples. **(B)** The cell trajectories labeled by the pseudotime analyzed by monocle2. **(C, D)** The circle plot of the strength of cell-to-cell communication in the MHCII signaling pathway network of glaucoma and normal samples, respectively. **(E, F)** The heatmap with the role of different immune cell types in the cell communication of the MHCII signaling pathway network of glaucoma and normal samples base on CellChat.

In the endothelial-fibroblast-pericyte cell cluster, ZFP36 exhibited the highest TRUE count and served as both a marker for this cell cluster and a switch gene involved in the transformation of fibroblasts into pericytes or endothelial cells. Additionally, it was found to affect heat shock protein binding activity and cellular response to cytokine stimulus. Furthermore, ZFP36 was identified as a DEG in the negatively associated module specific to POAG in the TM dataset. Notably, several genes with immune-related functions, such as CD74, DUSP1, and HLA-DPA1, which significantly affect the immune cell cluster, are also crucial in this endothelial-fibroblast-pericyte cell cluster. Furthermore, some biomarkers for neurodegenerative diseases (CSRNP1and SERPINA3) and genes associated with membrane integrity, barrier construction, extracellular matrix, and fibrosis (PTP4A1, RASD1, ADH1B, FMOD, MXRA8, PCOLCE, and SLPI) were also essential intersection genes. Changes in the expression of these genes affect the immune barrier. Complex changes in cell communication have also been observed in this cluster. Moreover, the MHCII pathway involving the HLA-D family (HLA-DPA1, HLA-DPB1, HLA-DQA1, HLA-DMA, and HLA-DRB5) and CD4 receptor-ligand pairs were also involved, similar to what was observed in the immune cell cluster. Additionally, several other intersection genes, including TNXB, CDH2, ADM, CCL3, HLA-C, CD74, ITGB5, and ITGAM, served as ligands or receptors for intercellular communication. These interactions could impact ECM receptors, cell-cell contact, and secreted signaling pathways involved in processes such as TENASCIN, CDH, CALCR, CCL, MHCI, MIF, VTN, APP, CD40, complement, CD23, GP1BA, ICAM, JAM, and THYI.

In the melanocyte cell cluster, numerous intersection genes emerged as both marker genes and participants in the transition from a normal to glaucoma state. Alongside the previously mentioned genes (DUSP1, RASD1, CEBPD, and ADM), the following genes exhibited high TRUE counts: LDLR, TYRP1, RPS12, DDIT4, and GADD45B.

## Discussion

As one of the most extensively studied sight-threaten ocular diseases, glaucoma often manifests as abnormal IOP. Unfortunately, despite significant research efforts, there is currently no standardized treatment available to effectively control IOP or slow the disease progression^3,7^. Additionally, the molecular mechanisms underlying the development of glaucoma remain poorly understood, primarily due to the complex nature of the IOP regulation system, which involves multiple tissue types and intricate microenvironments. In this study, we conducted a systematic molecular profiling of both POAG bulk microarray datasets from TM and single-cell RNA sequencing data from the choroid of AMD patients with glaucoma. Our objective was to elucidate potential pathomechanisms contributing to the glaucoma development. The TM, as the primary pathway for AH outflow, is expected to have the most direct and critical impact on IOP regulation. However, the choroid, which is part of the uveoscleral outflow pathway for AH, also plays an unneglectable role in IOP regulation, albeit being less explored in the context of glaucoma.

The possible pathomechanisms in the TM of POAG patients are summarized in **Figure 19**, which highlights the DEGs and the hub genes from two WGCNA modules (midnightblue and lightcyan) most strongly associated with POAG. Additionally, important genes enriched by functional analysis were also shown. Among the top five most significantly enriched pathways, two are from morphological alterations of TMCs and the ECM imbalance pathways. These changes have been widely discussed in previous research papers ^12,63^. The dysregulation of cell morphology and its functional imbalance mutually reinforces each other. Together with ECM disorder pathway, characterized by excessive matrix deposition, these alterations result in decreased TM elasticity and narrowing of the AH outflow channel, causing the obstruction in the flowing of ocular fluid ^14^. Similar findings were highlighted in the glaucomatous choroid, where pathways enriched in the endothelial cell-fibroblast-pericyte cluster were concentrated on ECM and actin-binding activities during the transition from pericytes to fibroblasts. Taking all together, our work suggest thickening of the vascular wall has a direct effect on the IOP regulation ^58,64^.

**Figure 19:**
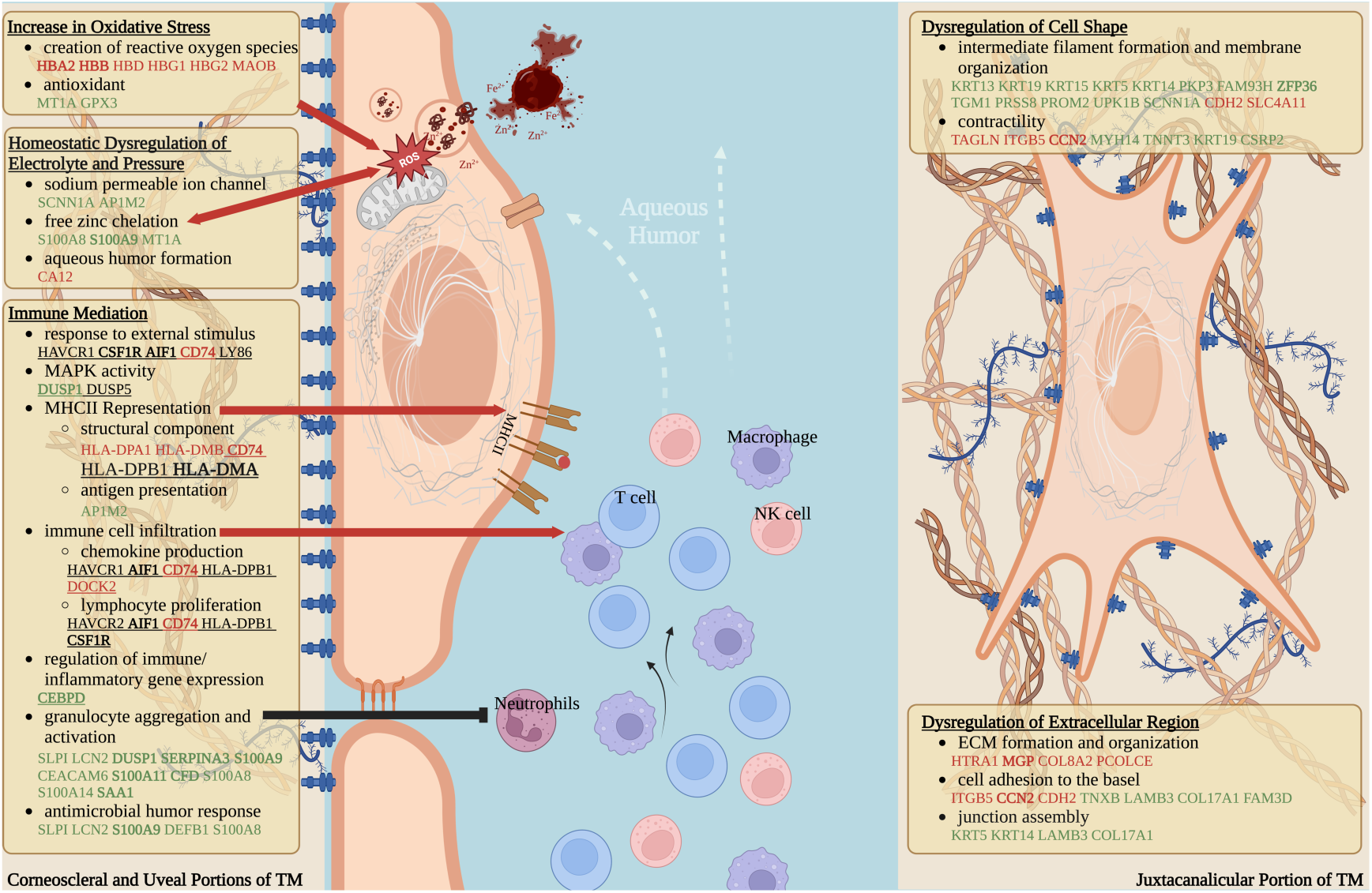
Suspected Pathological Changes in TM of POAG. Genes color-coded in red are upregulated, in green are downregulated, and in black are stable in POAG as compared to the control group (cut-off: *adj.P* < 0.05, | log2FC | > 1). Genes with underline indicate that they are hub genes from the POAG-related modules assessed by WGCNA. Genes in bold were hub genes of DEGs.

Next, we systematically examine the most differentially expressed genes in glaucoma patients compared to control. The DEGs with the highest degree of differential expression in the TM datasets compared to control were HBB, HBD, and HBA2, all of which are hemoglobin components. According to histopathological studies, the nutrients and oxygen of the TM are supplied by the AH flowing through it rather than through the blood vessel ^15^. Therefore, we hypothesize that the increased expression of hemoglobin-related genes could be the result of hemolysis, and the increased expression may have come from platelet-derived microparticles. We further speculated that an elevation of hemoglobin-related genes may lead to an increase in iron and zinc ions ^65^. In line with this hypothesis, studies have reported elevated levels of zinc and iron ions in the AH of glaucoma patients, and laboratory experiments have validated similar results ^66,67^. The increased levels of free hemoglobin fragments and free metal ions are essential sources of ROS and cause serious damage to mitochondria ^68,69^. In addition, some antioxidant-related genes like MT1A and GPX3 were found to be downregulated (refer to **Fig. 19**), which further explained the fact that oxidative stress was significantly enriched in the functional analysis. Notably, genes involved in zinc ion chelation, such as S100A8, S100A9, and MT1A, were also downregulated in this study (see **Fig. 19**), potentially impacting the neutralization of excess zinc ions and contributing to oxidative stress. Excessive oxidative stress can lead to mitochondrial structural disruption and affect oxidative phosphorylation ^68^, which aligns with the findings of this study.

Interestingly, oxidative stress-induced immune alterations have been increasingly recognized as important factors in glaucoma development. The immune system’s role in glaucoma, similar to other neurodegenerative diseases, is not yet fully understood, and effective biomarkers for disease progression and chronic inflammation are lacking ^70^. Therefore, this study focused on investigating immune alterations in the glaucomatous microenvironments and found significant abnormalities:

Multiple DEGs with immune or inflammatory functions were identified in both TM and choroid data, and the intersecting genes with the highest TRUE counts were immune-related. In addition, the enrichment of the MHCII pathway, indicating potential activation of CD4^+^ T cell-mediated immune response ^71^, was particularly evident. Consistently, infiltration of non-CD8^+^ T cells was identified in the TM of POAG samples, and both top POAG-related WGCNA modules had the closest relationship with T cell infiltration. Similar enrichment of the MHCII pathway was observed in the choroid of glaucoma samples, with NKT cells being significantly enriched as influencers. These results were in line with the findings of increased T cell-mediated immunity in the peripheral blood of glaucoma patients and the retina of glaucomatous animal models ^72–75^, indicating a similar important role of T cell immunity in both the glaucomatous TM and choroid. hTM cells can express class II antigens of the MHC ^76^, while macrophages, which were found to infiltrate glaucomatous TM in this study, were also able to trigger T cell aggregation and activation. Therefore, a more precise study, such as a single-cell transcriptome analysis of glaucomatous TM, is needed to distinguish the specific cells involved in activated immunity.

The switch from NKT cells to macrophages observed in the immune cell cluster of the choroid single-cell transcriptome data is intriguing. Previous studies have highlighted the importance of retinal glia, including microglia, astrocytes, and Müller cells, in the initiation and progression of glaucoma ^77^. Microglia have been shown to trigger immune-inflammatory T-cell infiltration into the inner side of the blood-retina barrier, although the exact patterns of blood-retina barrier impairment are yet to be fully understood ^78,79^. As the retina, especially the optic nerve head, is the direct and possibly initial site of glaucomatous damage ^77^, it is inferred that activated T cells within the blood-retina barrier cross back to its outer side to the choroid, influencing the infiltration of macrophages and exacerbating immune abnormalities in the choroid microenvironment. This suggests a complex interplay between various immune cell types and glial cells in the pathogenesis of glaucoma. Further investigations that consider changes occurring both within and outside the blood-retina barrier, are necessary to elucidate the specific mechanisms and interactions involved in this process.

## Conclusion

By integrating bulk microarray data from the TM and single-cell transcriptomic data from the choroid, we were able to examine and correlate glaucomatous abnormalities between the anterior and posterior ocular tissues involved in IOP regulation. Both studies showed immune alterations, particularly the enrichment of the MHCII pathway and T-cell infiltration, hold promise for the development of biomarkers and targeted treatments for glaucoma.

However, it should be noted that the specific cell types within the TM that contributed to these alterations could not be distinguished solely from the bulk microarray data. Moreover, limited by the availability of public data, the single-cell transcriptome data from the choroid utilized were from AMD patients, and the subtypes of glaucoma cases were not clarified. Therefore, it is crucial to conduct further experiments that investigate the specific cell types and corresponding molecular changes associated with different subtypes of glaucoma by single-cell RNA-sequencing or other platforms. This will provide a more comprehensive understanding of the disease and facilitate the realization of the aforementioned goals, including the identification of biomarkers and the development of targeted treatments for glaucoma.

