## Supplementary figures and images for "Integrative Transcriptomic Analysis of Anterior and Posterior IOP-Controlling Tissues in Glaucoma Reveals Enrichment of MHC-II Pathway and T-Cell Infiltration Signatures"

### fig2.tif

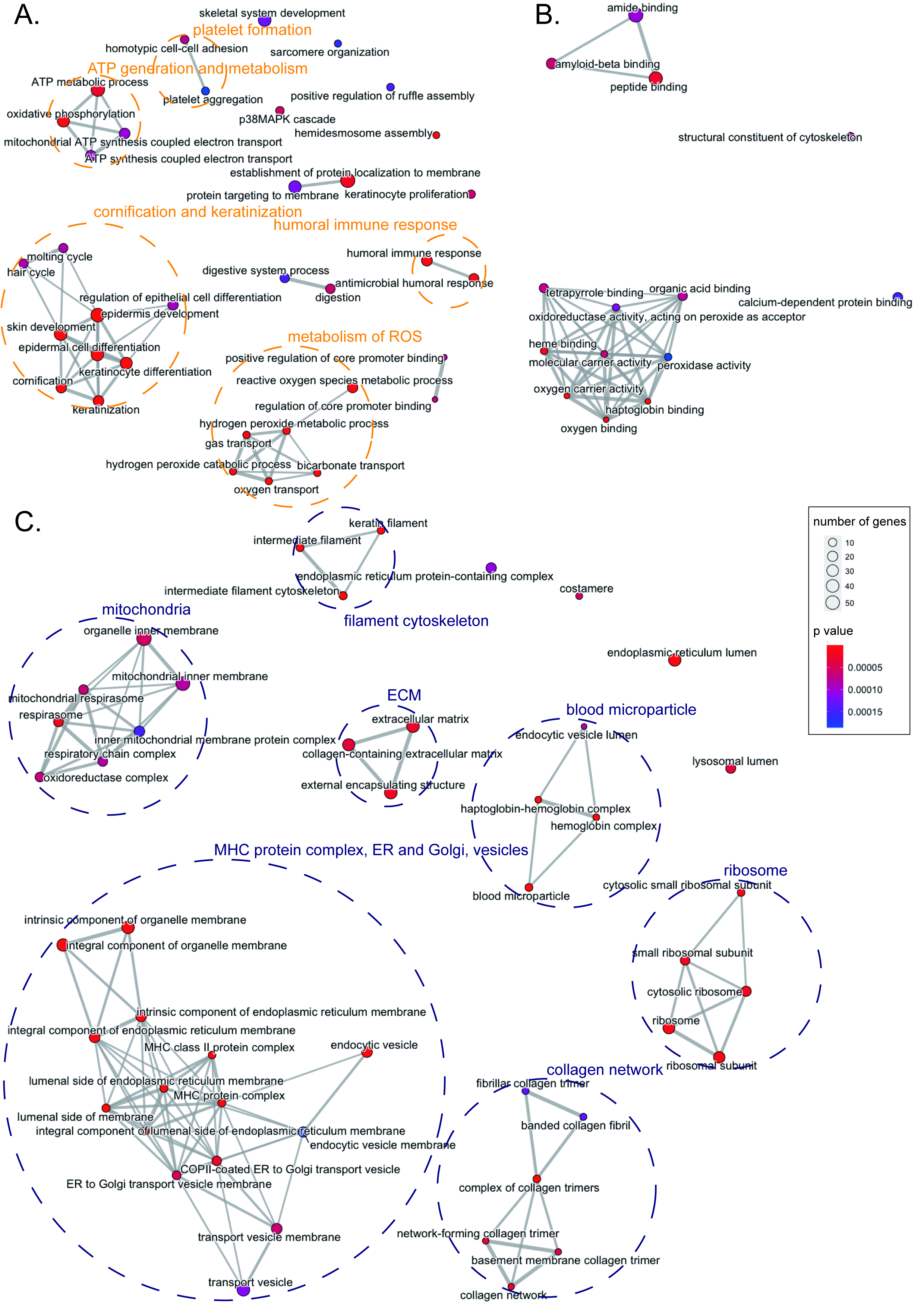

### fig3.tif

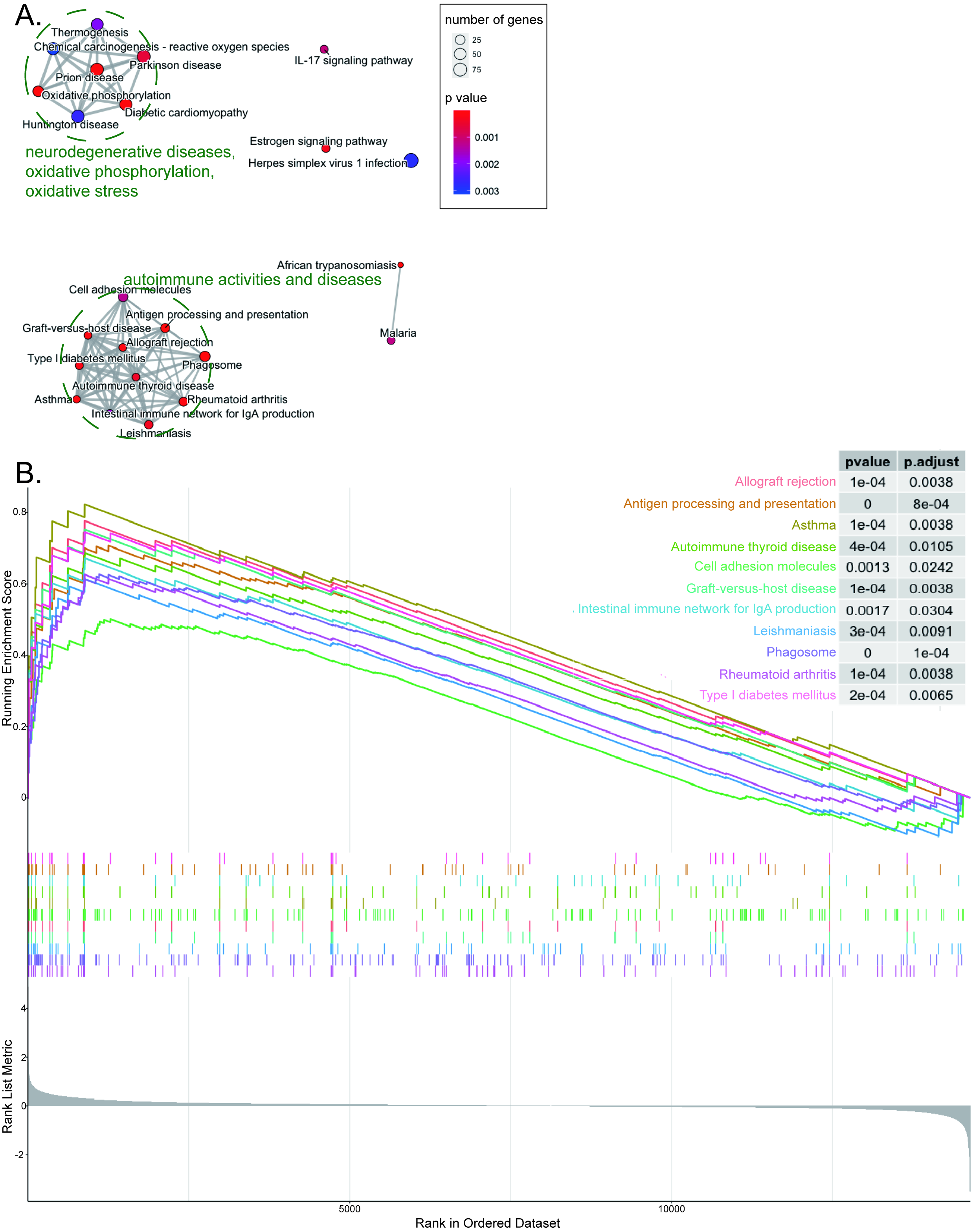

### fig4.tif

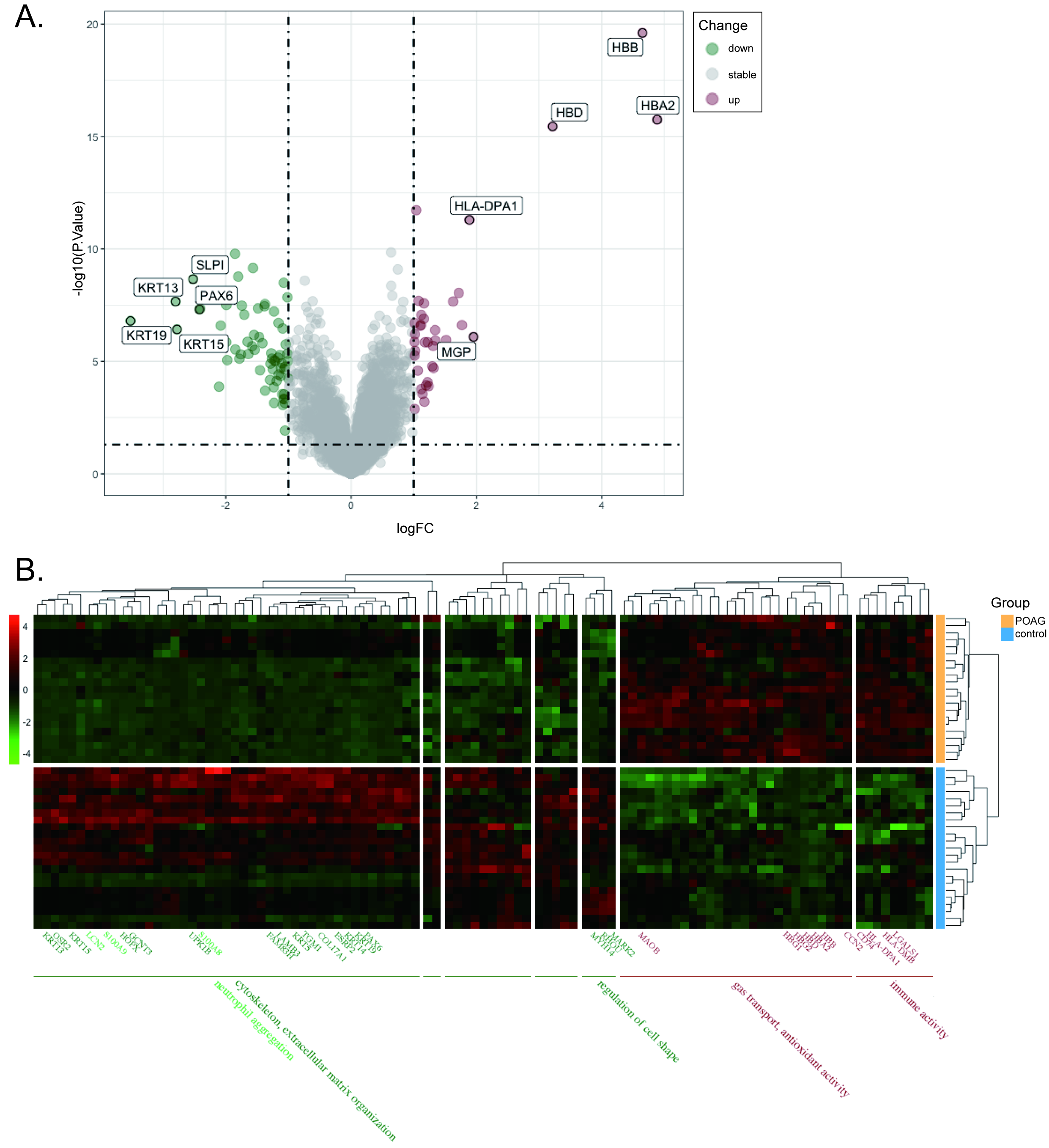

### fig5.tif

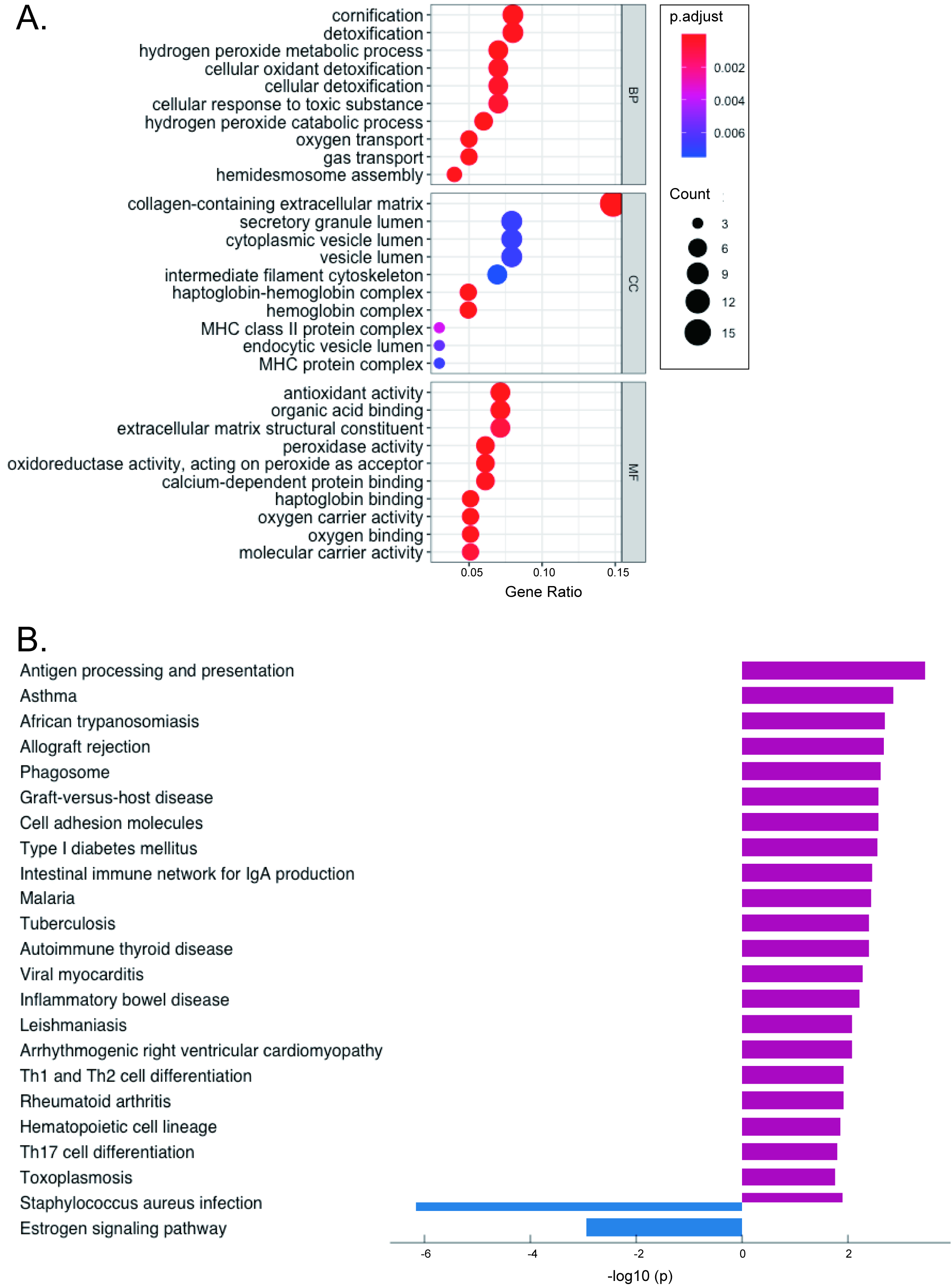

### fig6.tif

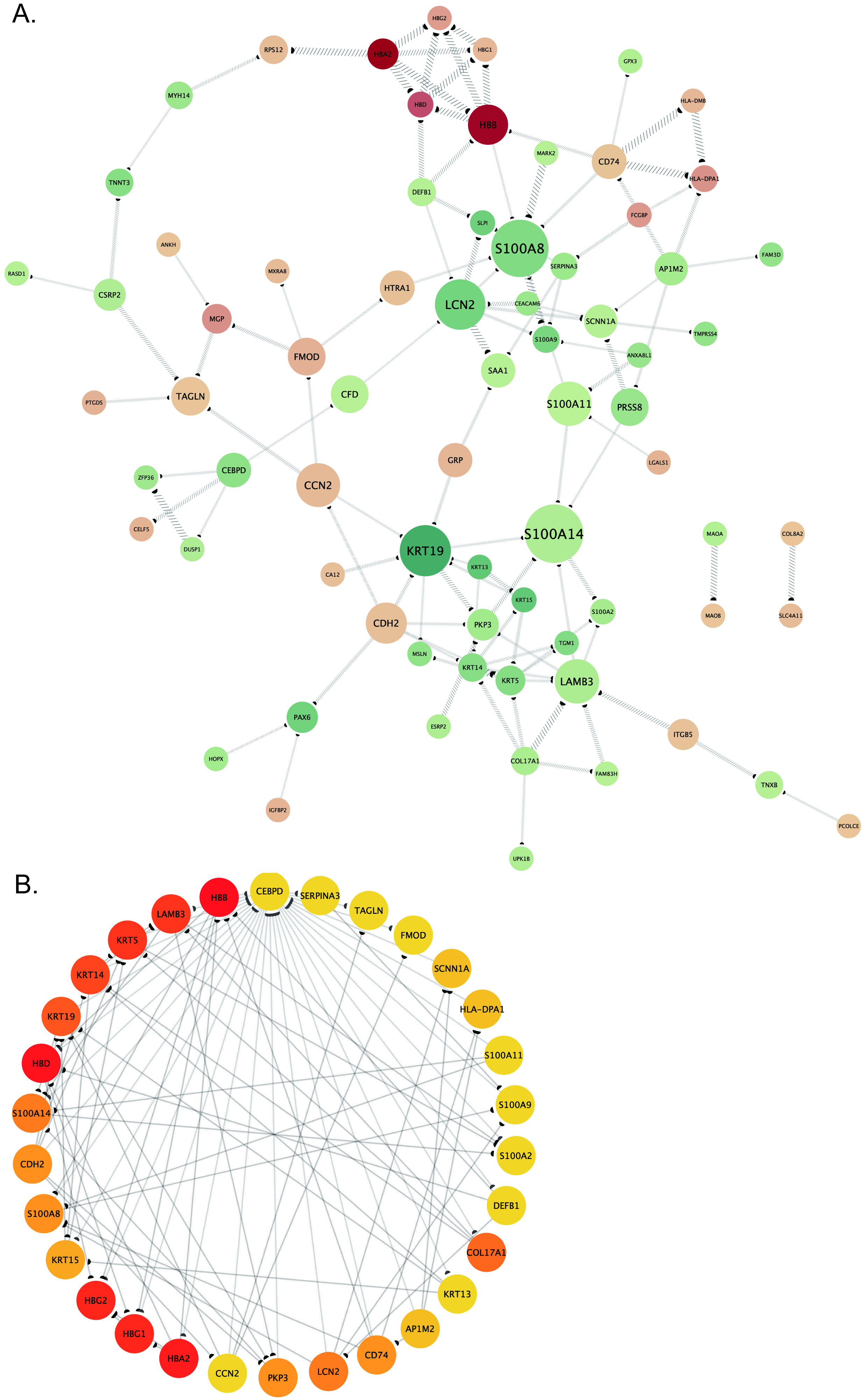

### fig7.tiff

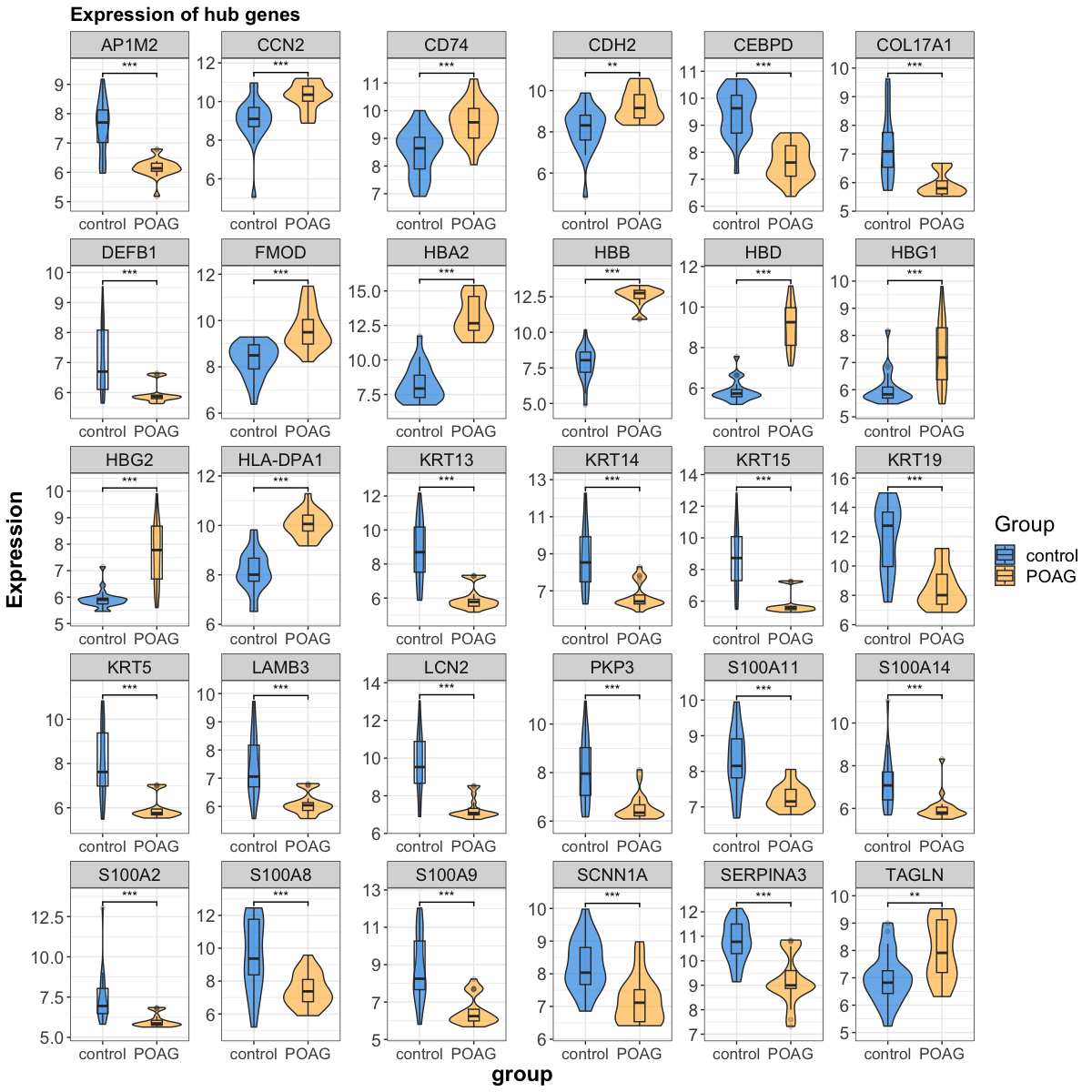

### fig8.tiff

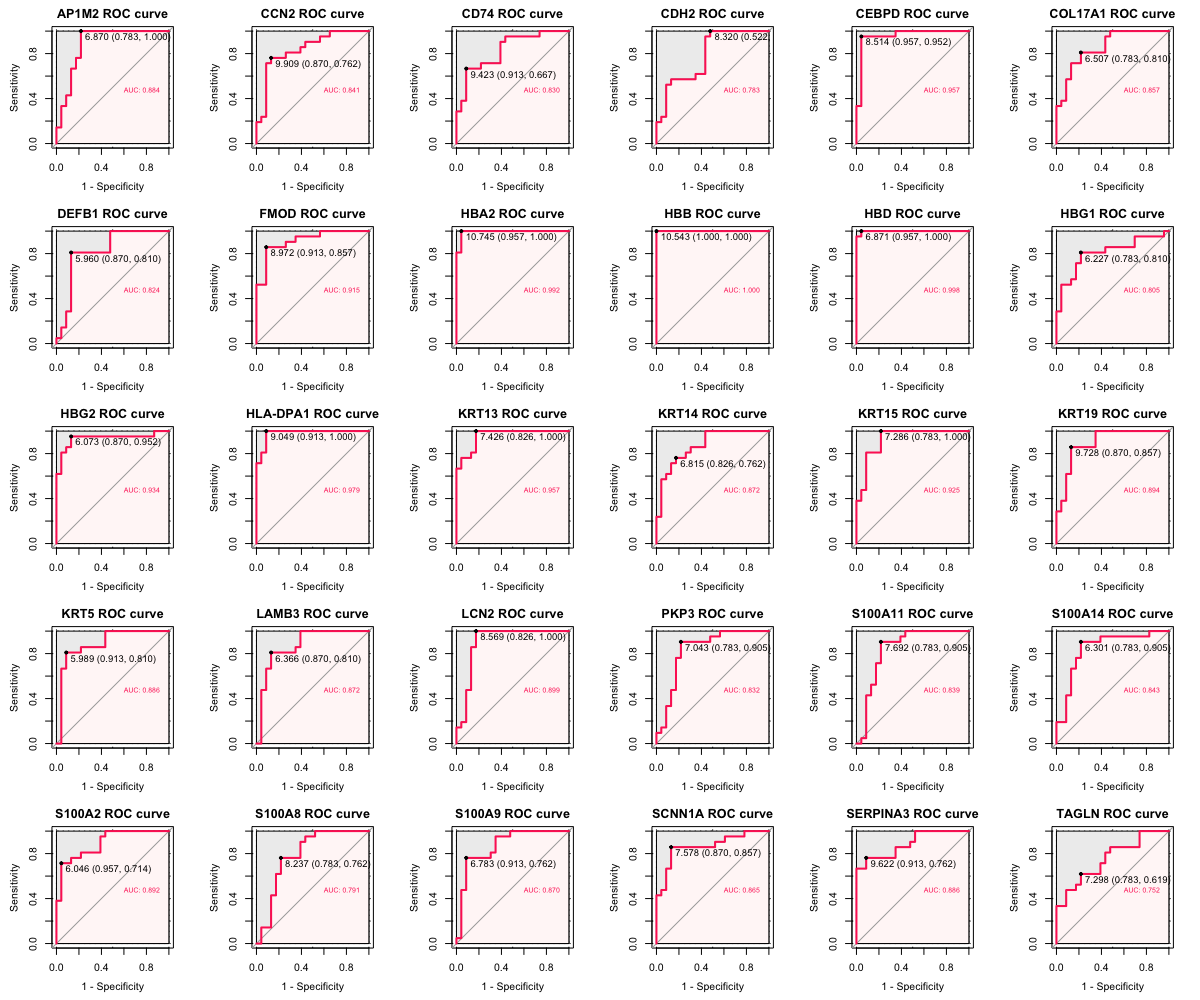

### fig9.tif

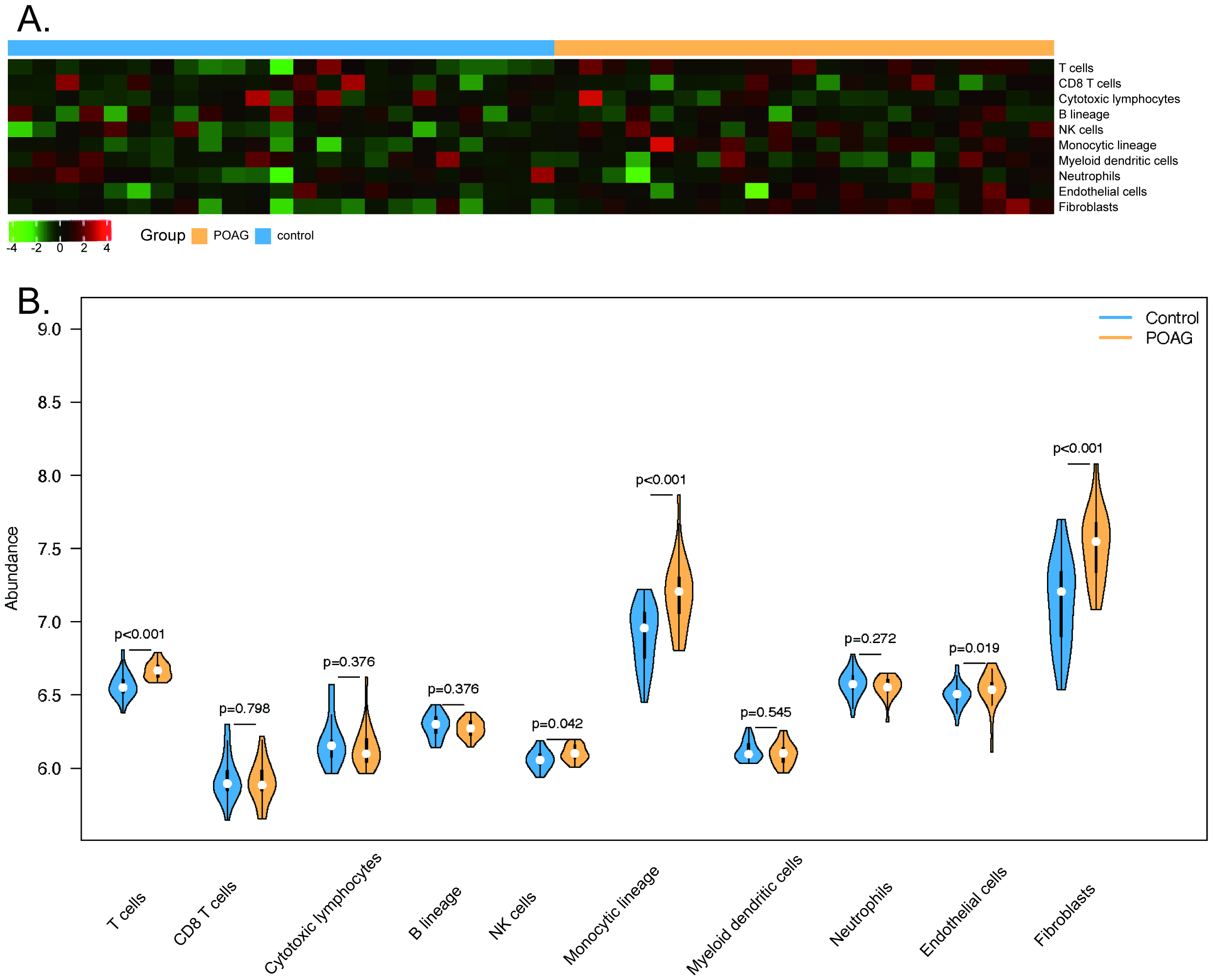

### fig10.tif

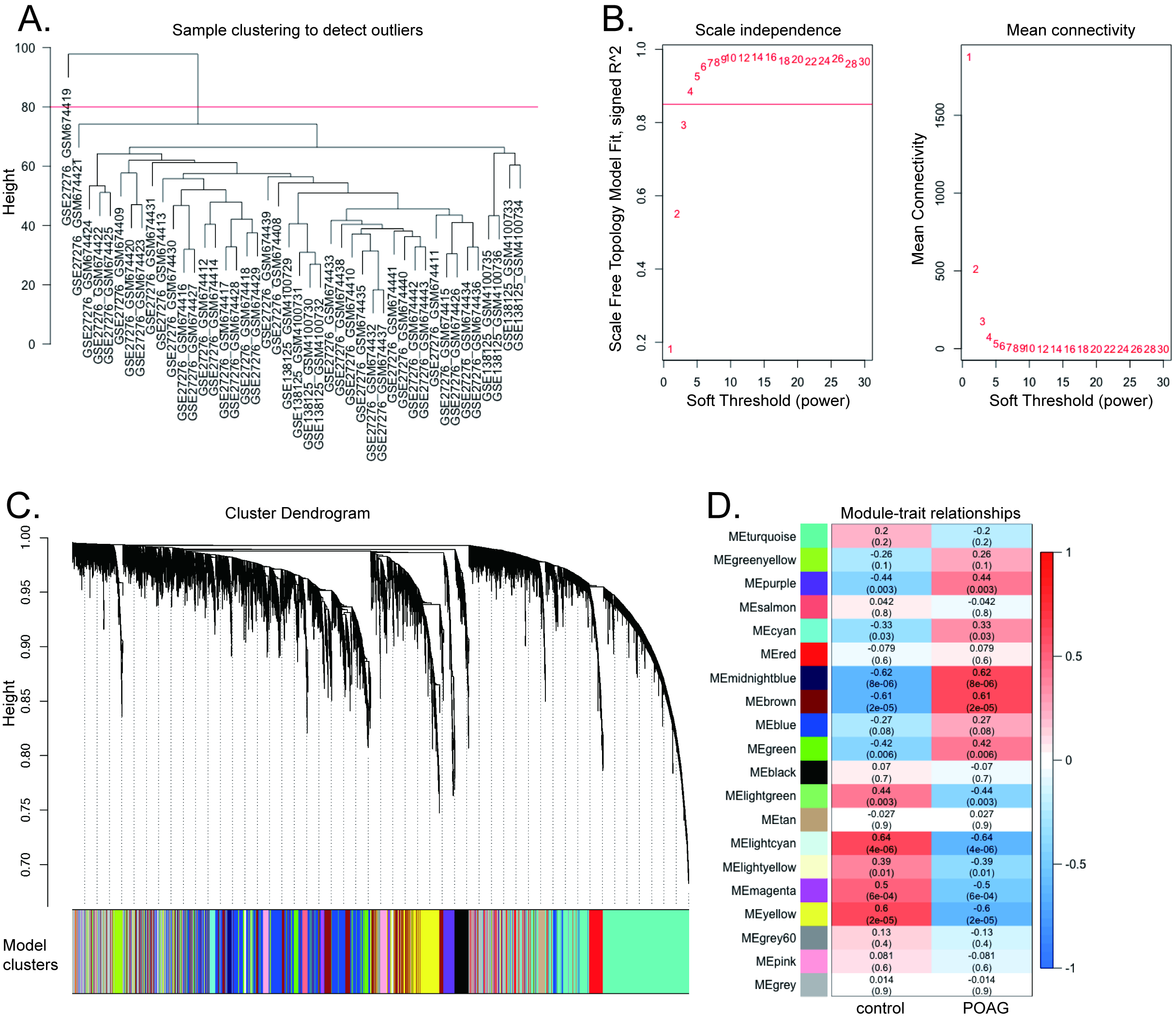

### fig11.tiff

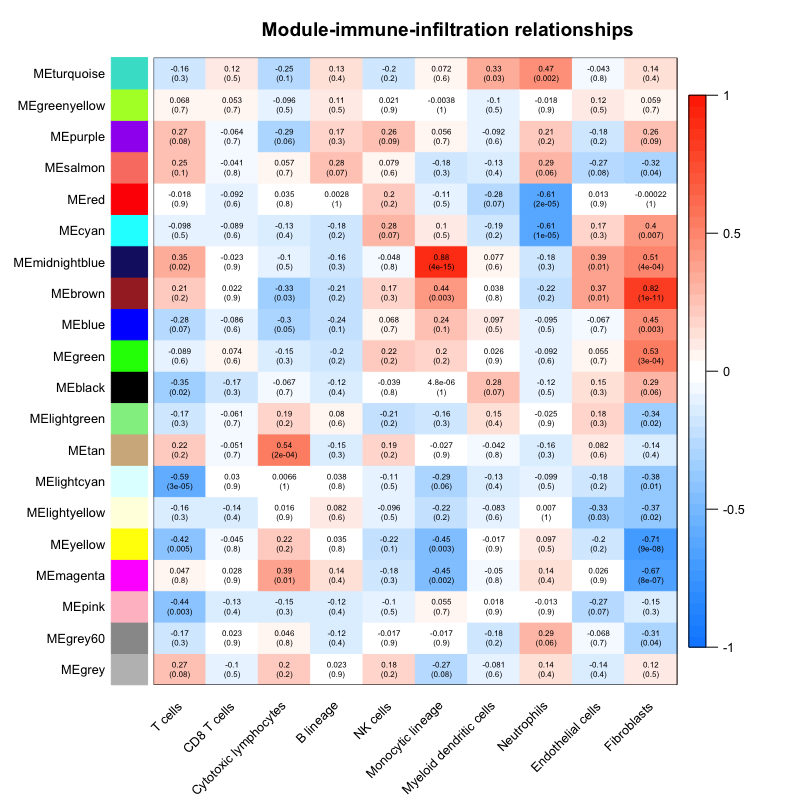

### fig12.tif

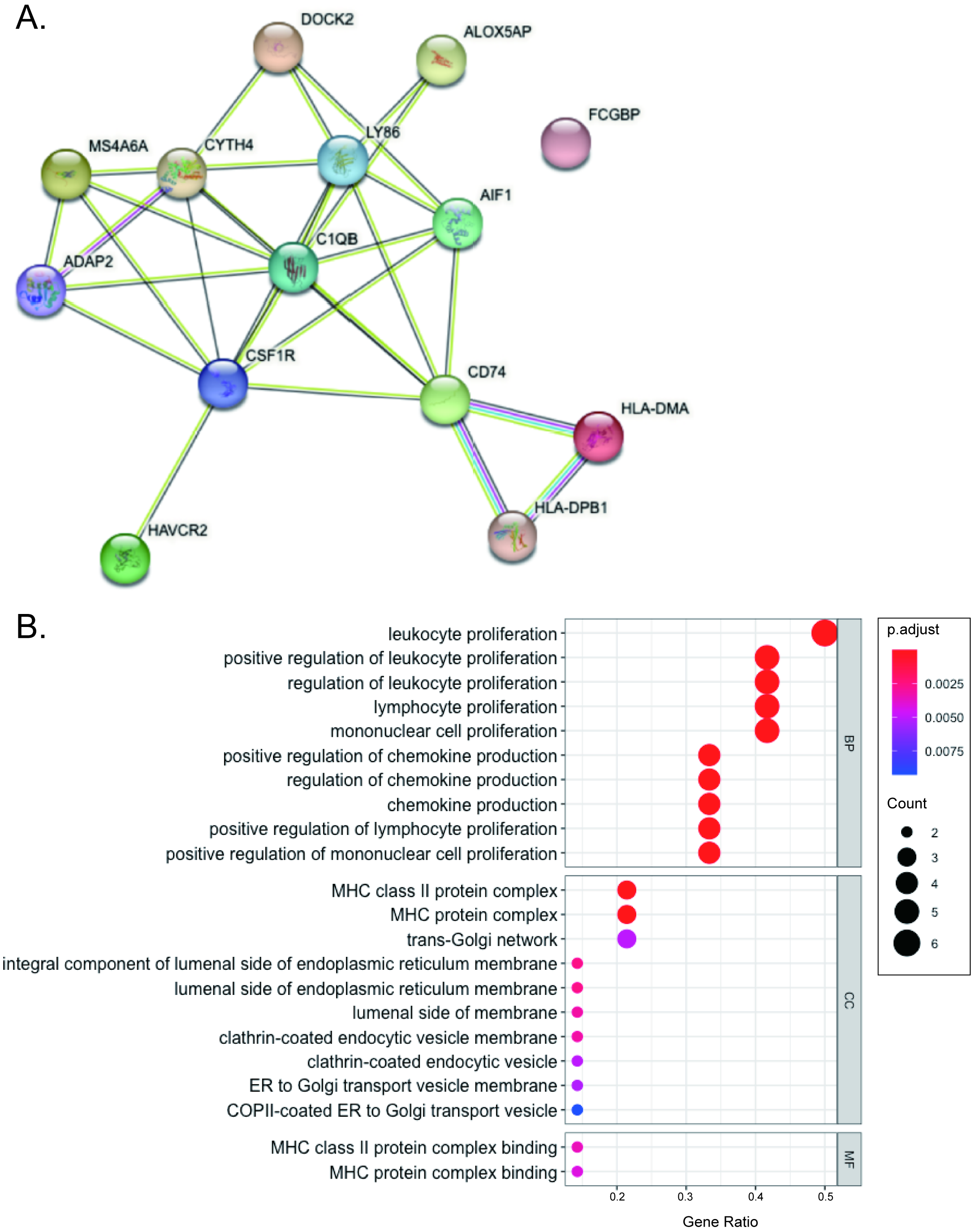

### fig13.tif

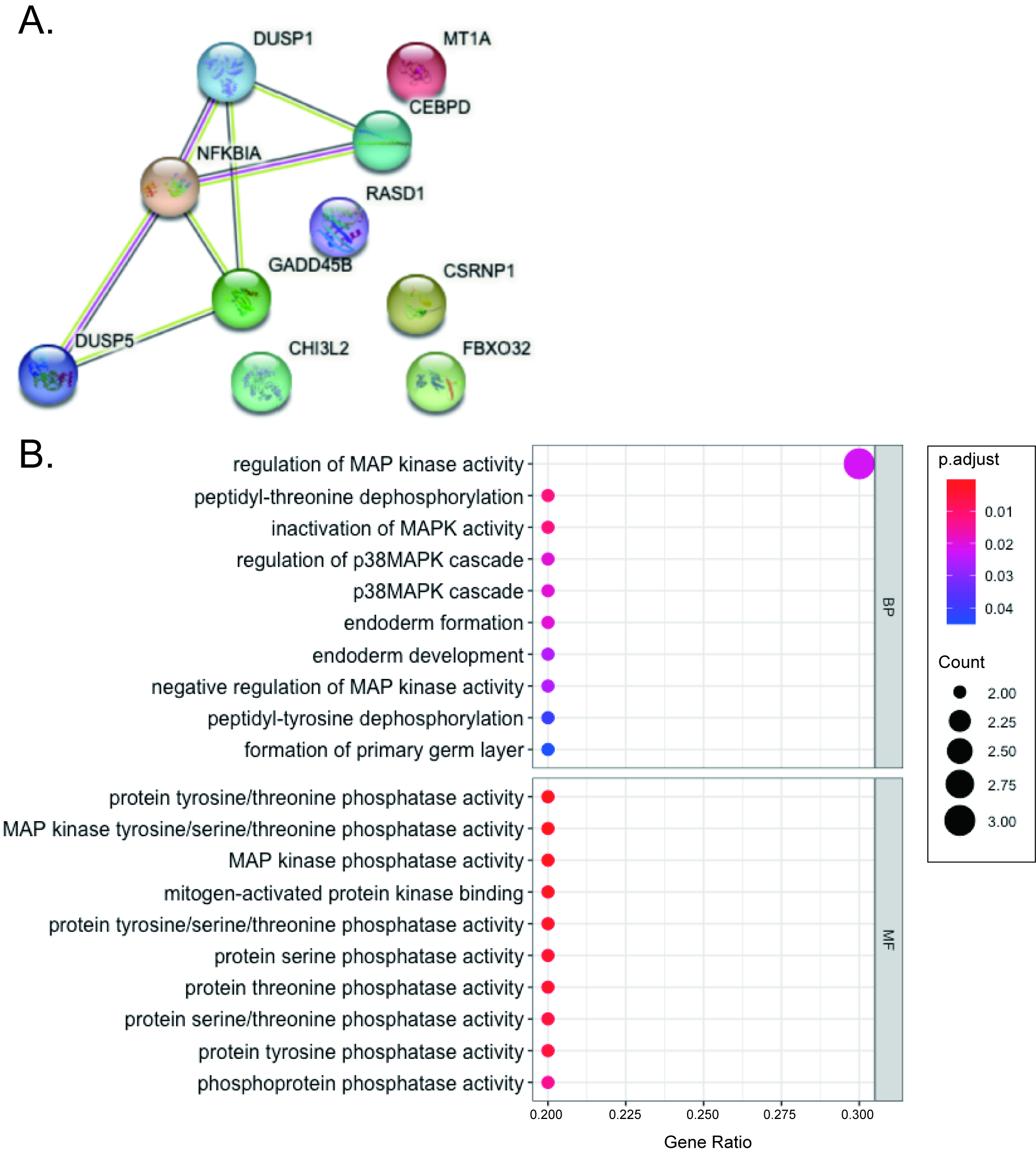

### fig14.tif

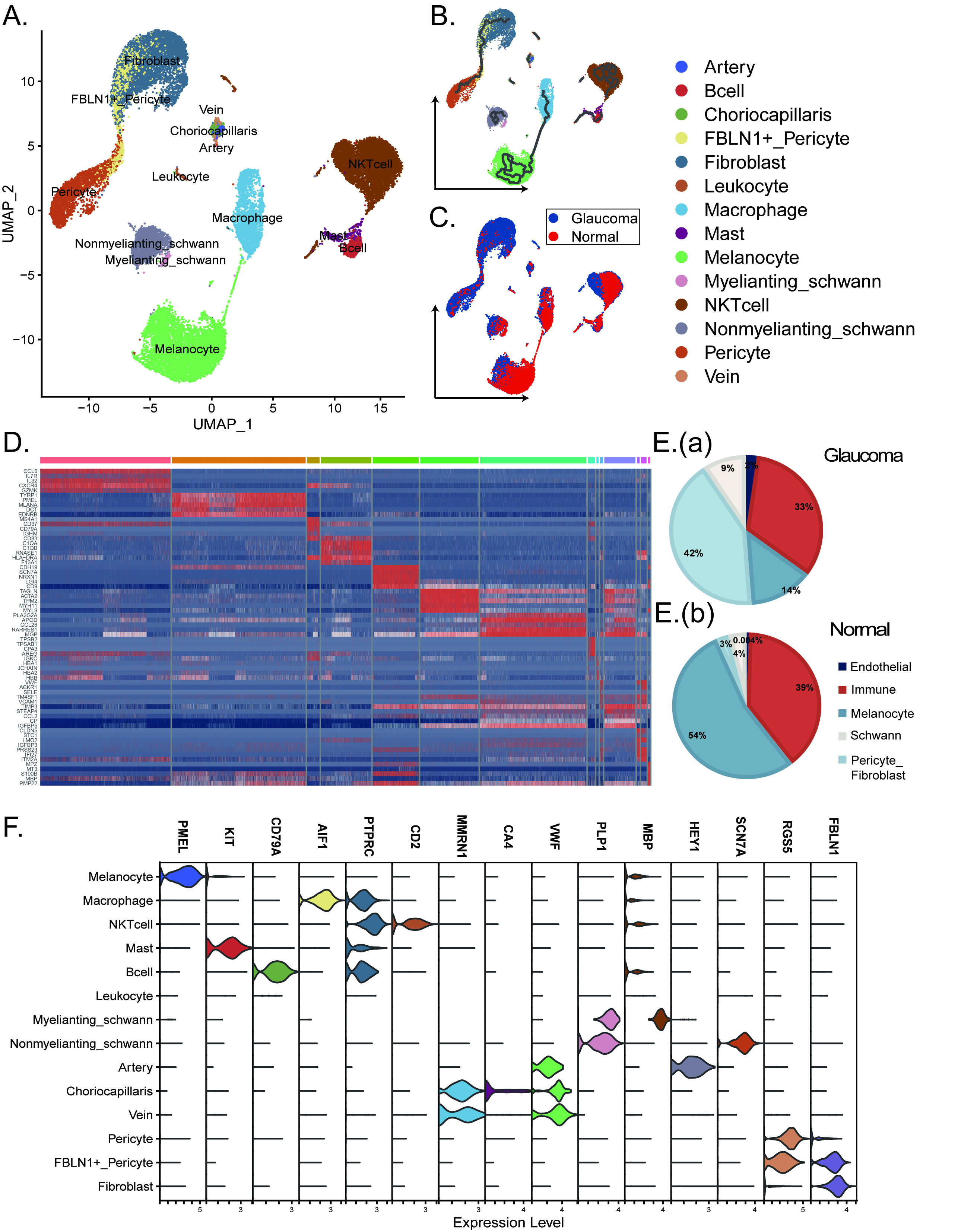

### fig15.tif

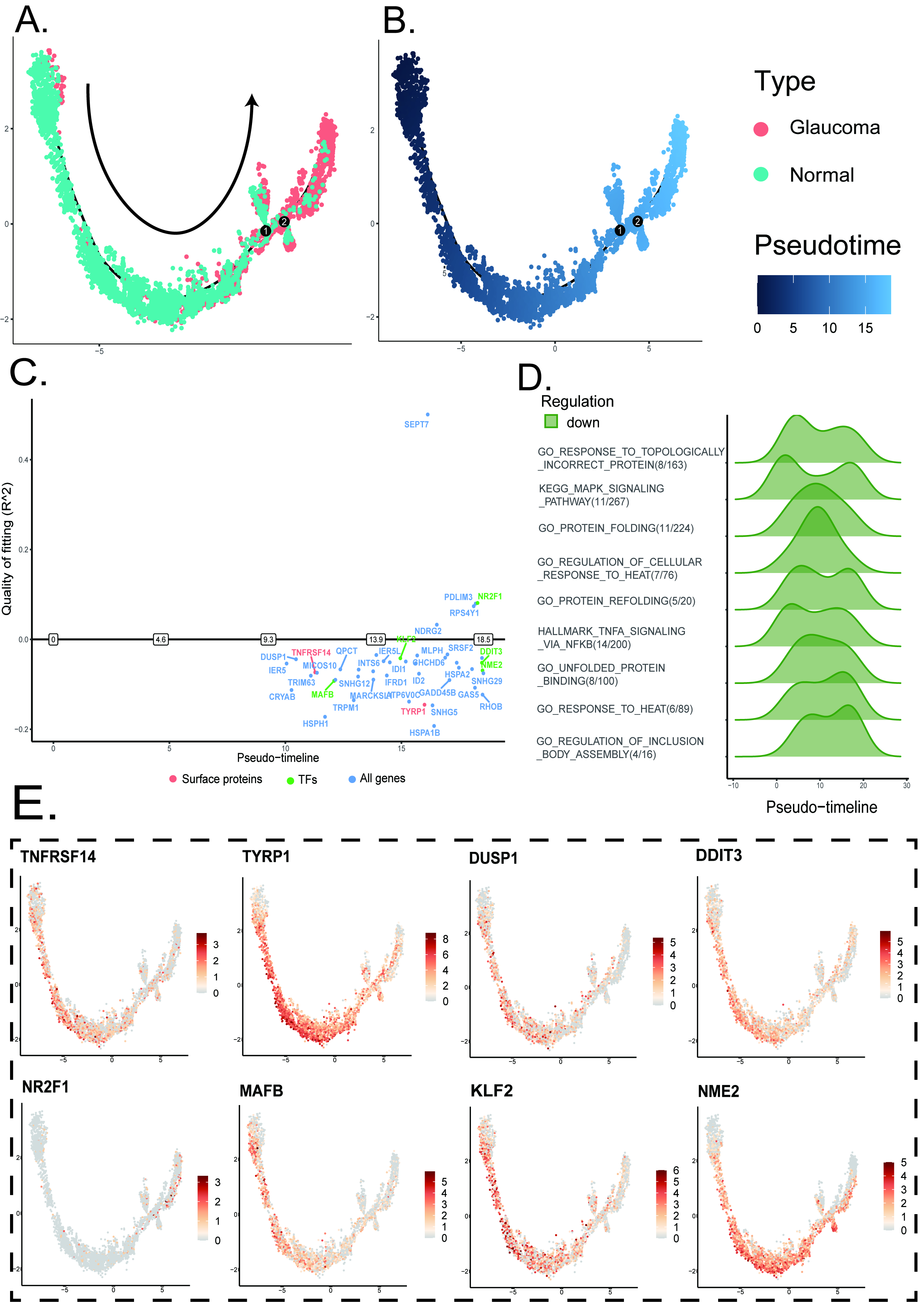

### fig16.tif

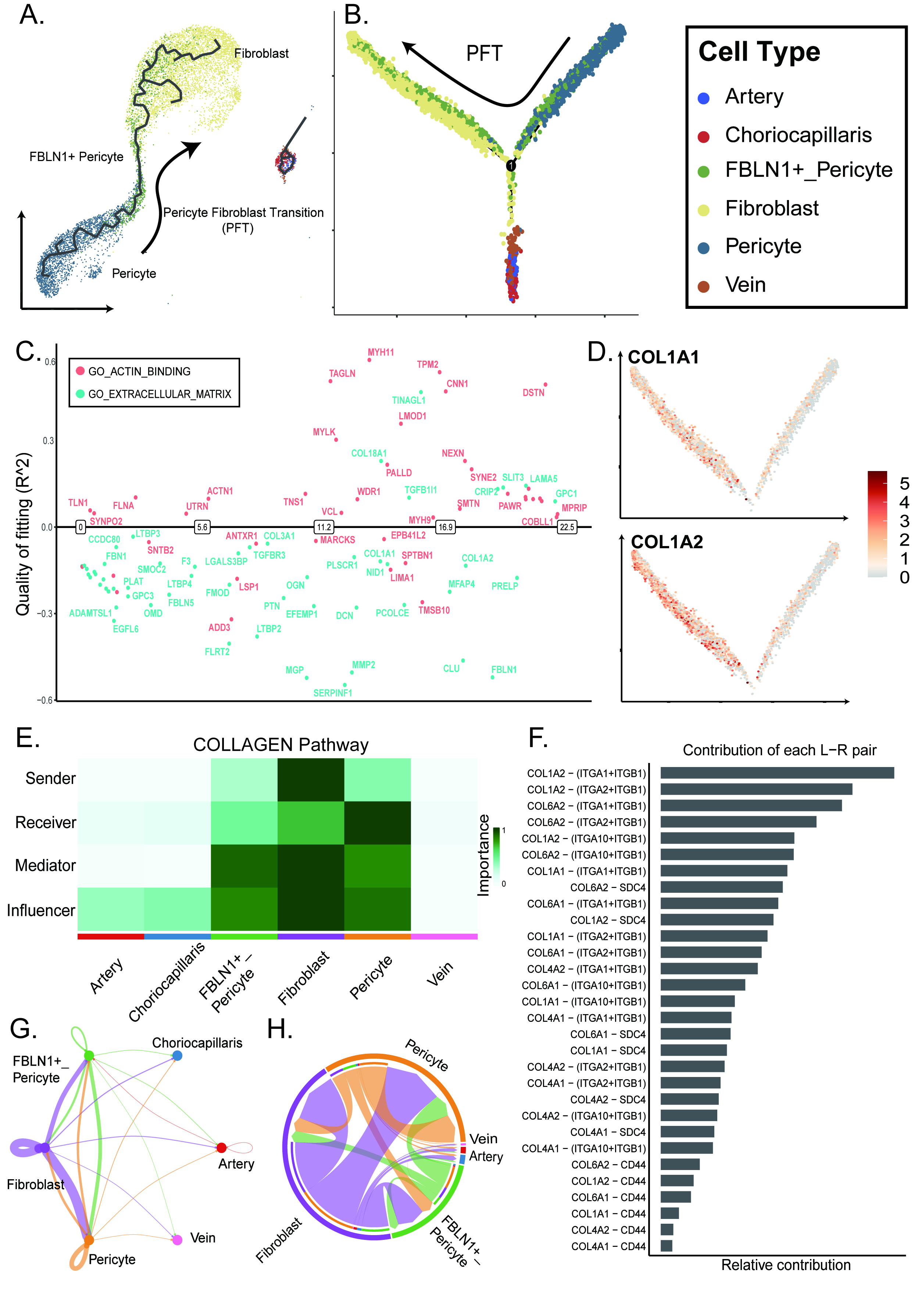

### fig17.tif

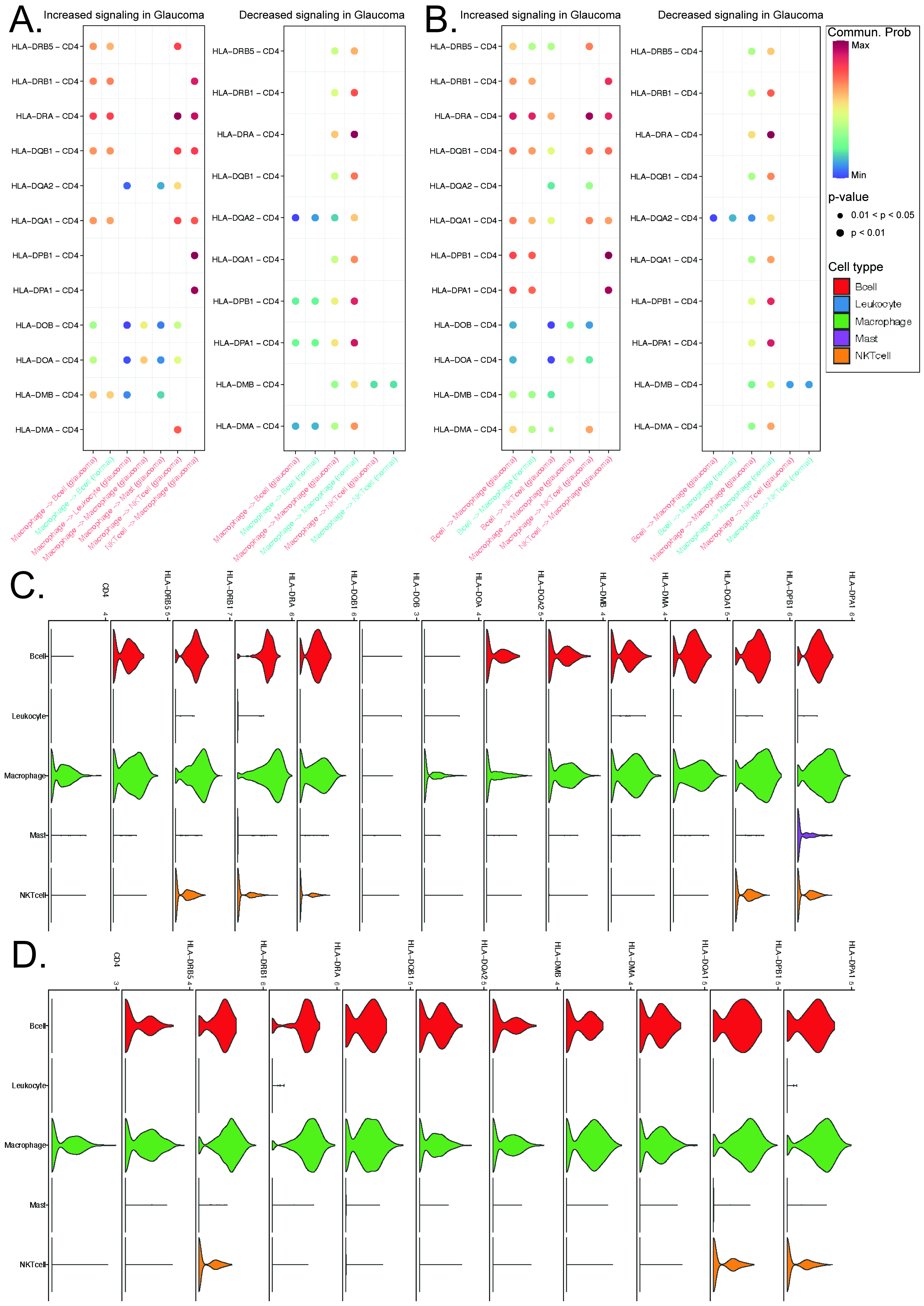

### fig18.tif

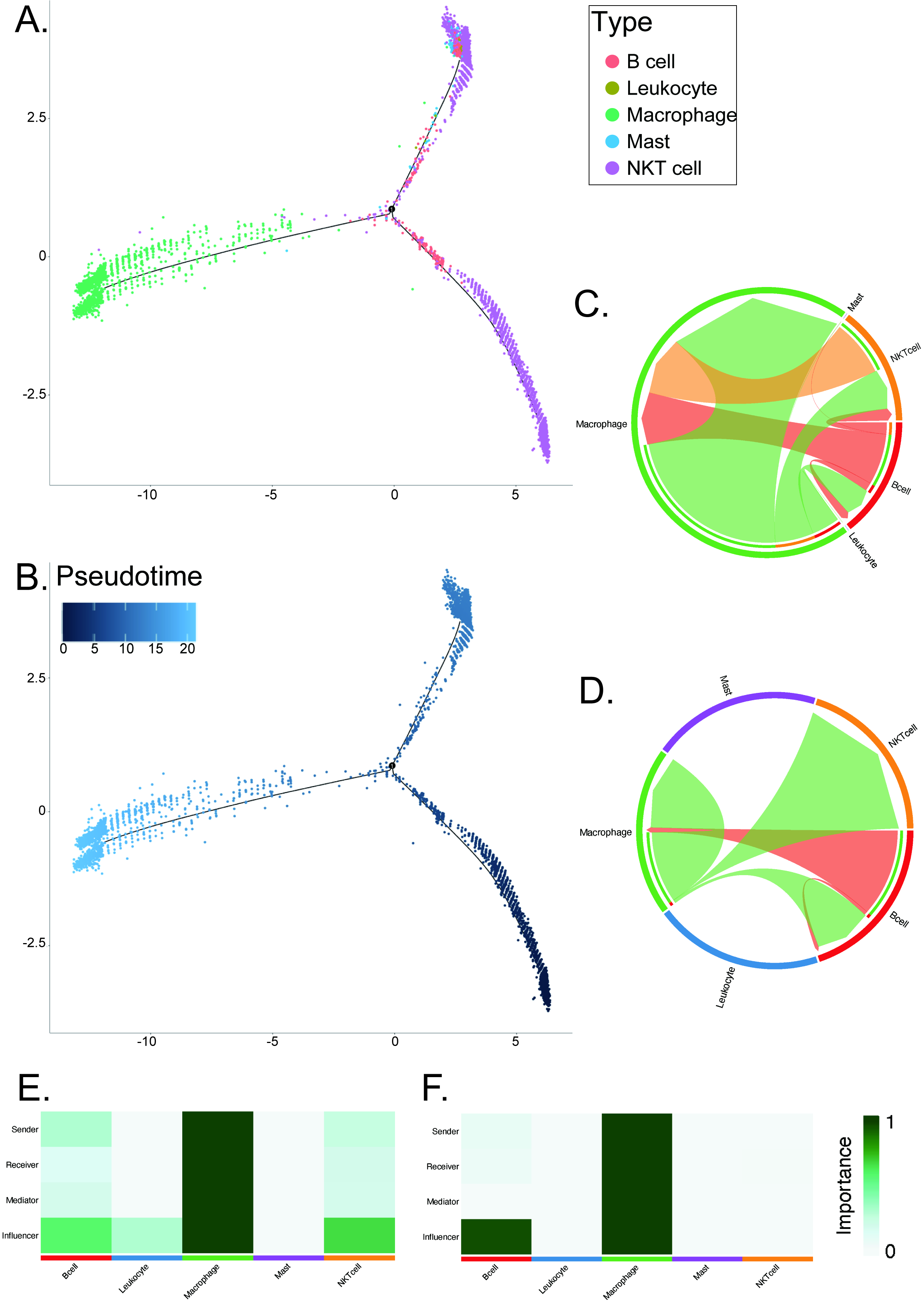

### fig19.tif

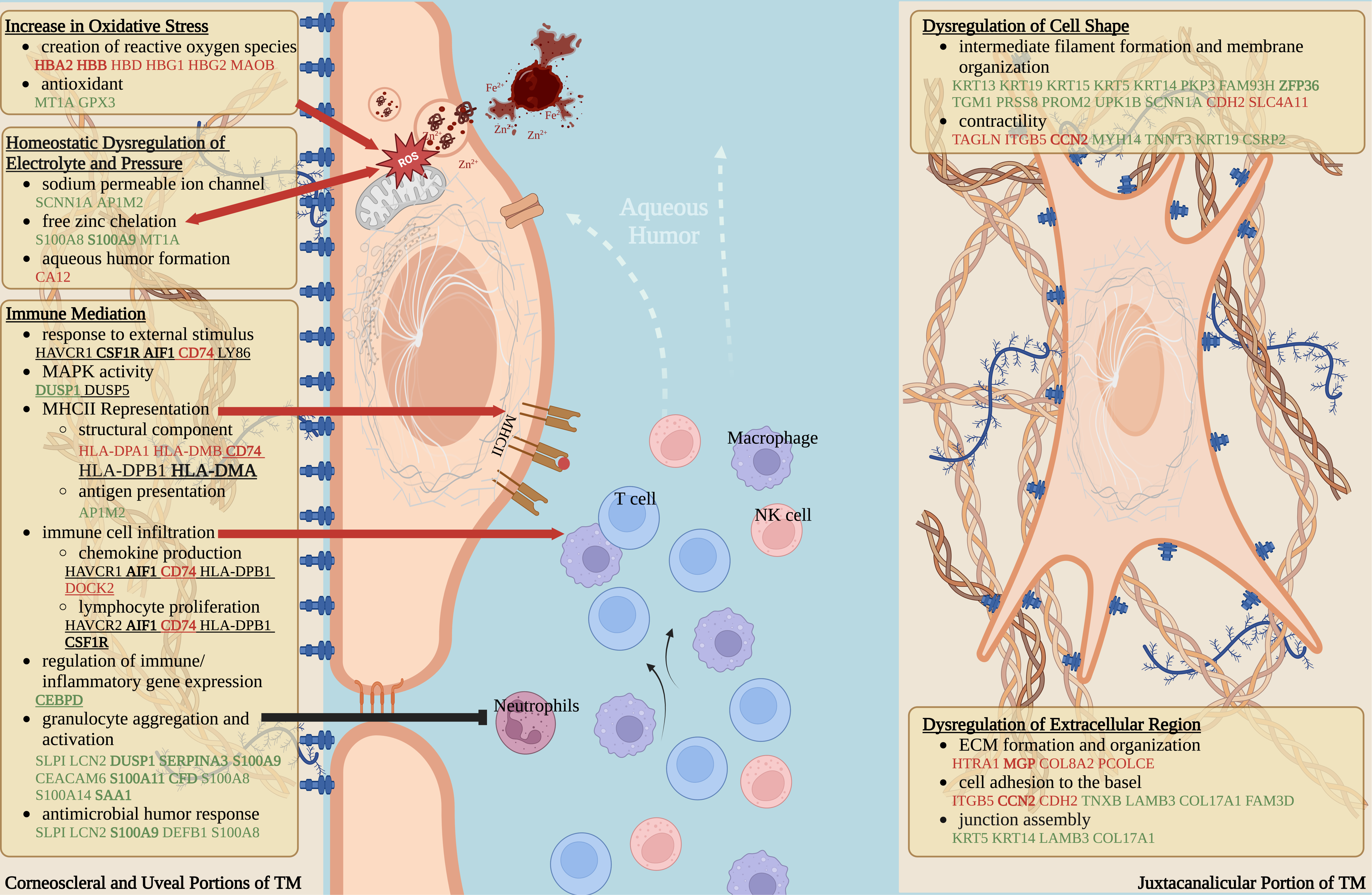

### suppl_fig1.tif

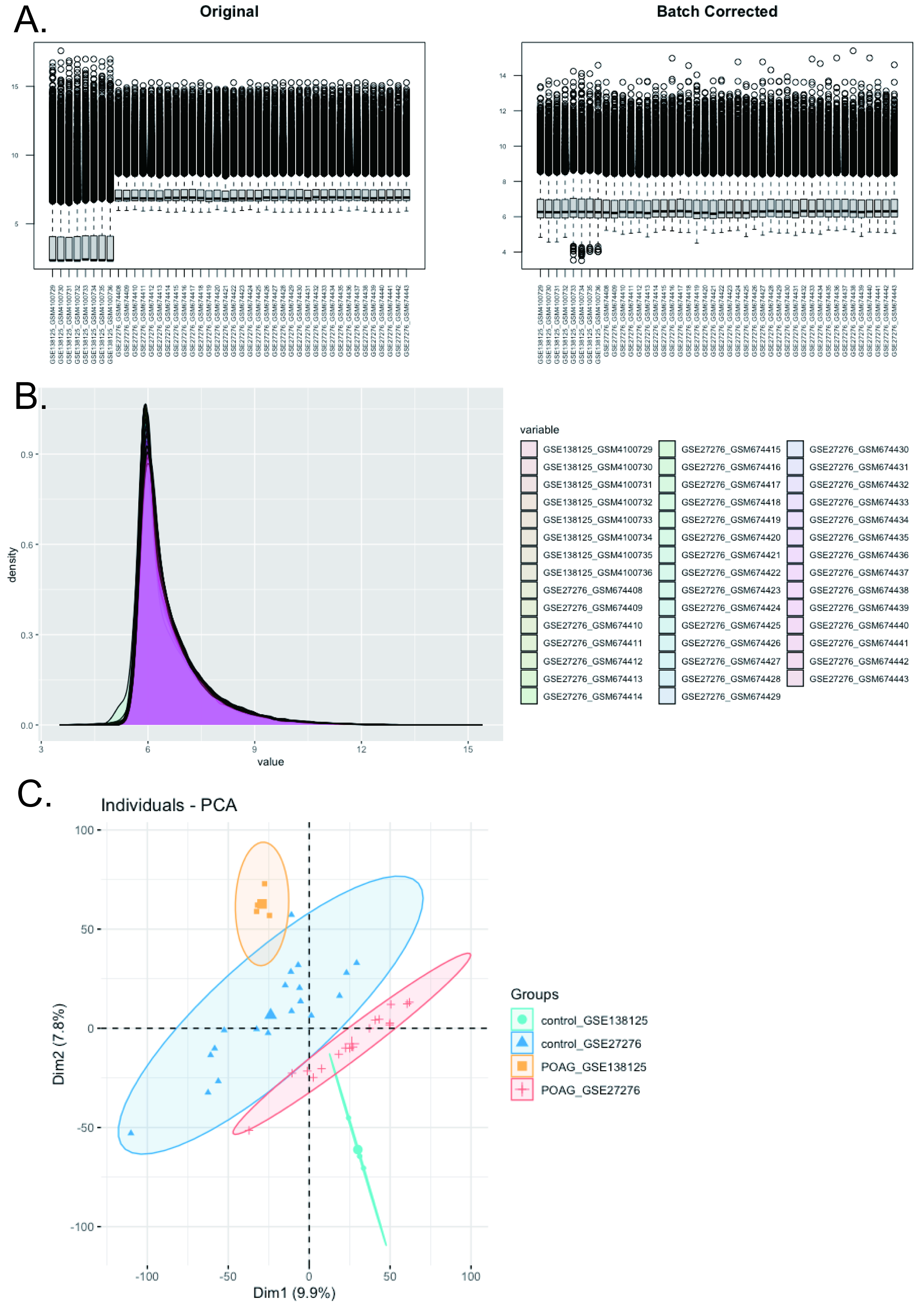

### suppl_fig2.tif

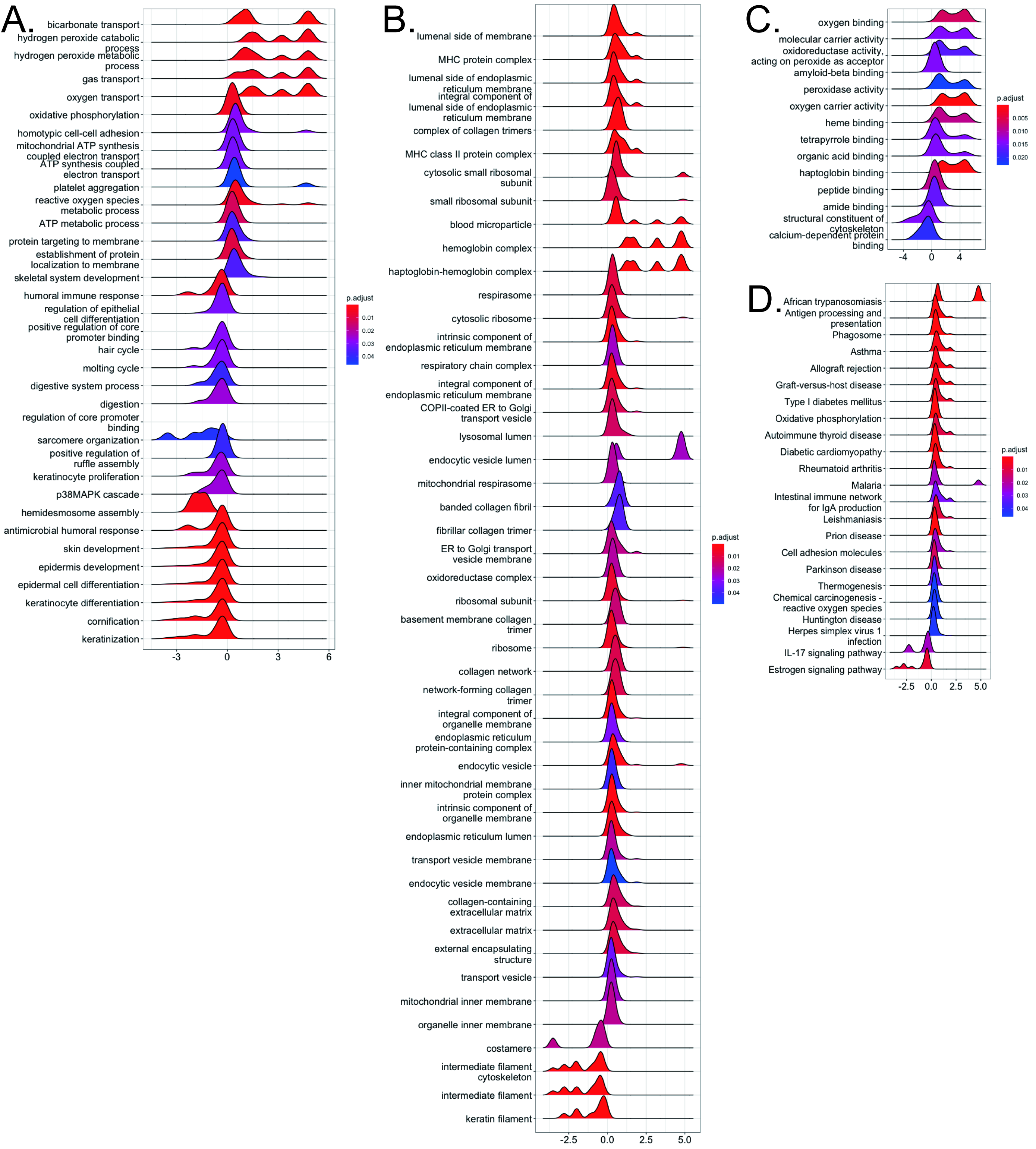
